# An atlas of human histological diversity

**DOI:** 10.64898/2026.01.30.702249

**Authors:** Hugo Mantion, Zhao Zhang, Diego Serra, Lætitia Lebrun, Maxime Tarabichi, Vincent Detours

## Abstract

Descriptions of tissues by histopathologists rest on verbal statements with limited quantitative accuracies. Here we propose an unsupervised computational framework that transposes to histology the concepts of RNA-seq gene expression analysis, setting histology on a quantitative ground. Leveraging this framework and the GTEx dataset, we built an atlas surveying the histological diversity of 40 organs from 946 non-diseased individuals and documenting 11,125,747 associations between 2,560 morphemes—the histological operational analogs of genes in our framework—and 9 layers of patho-clinical and multi-omic molecular data, providing a rich context to interpret histology. In contrast with the ideally healthy normal specimens depicted in histology textbooks, the atlas reveals the influence of age, sex, genetics and sub-clinical pathologies on tissue structures. For example, we report that female eccrine sweat glands are surrounded by more adipocytes than their male counterparts, and that distinct calcification-associated aorta morphemes are driven by either smoking or genetic polymorphisms. Cross-organ analyses also delineate the systemic histological impact of diabetes and other conditions and establish the power of blood gene expression to predict disease-related morphemes. The atlas is released as an interactive research tool to explore human histological diversity.

## Introduction

While histopathology is based on rigorous tissue observations, descriptions of morphological concepts typically rest on natural language statements. For example, the textbook description of thyroid cancer cell nuclei state that “*[cancer cell] nuclei have been variously described as pale, clear, optically clear, watery, empty, ground glass. […] Nuclei with these alterations are enlarged […]*”^1^. But what is the quantitative limit between a normal and an enlarged nucleus? Do two pathologists always agree on what ‘ground glass’ means? Such verbal descriptions contribute to the inter-observer variability recognized in a several diagnostic tasks^2^, and limits the scientific understanding of tissue morphologies. Artificial intelligence (AI) models trained with human supervision have the potential to improve on human diagnostic accuracy^3^, and expand its scope, for example by predicting cancer driving mutations^4^. But supervised training bounds scientific investigations within the limits of known categories and concepts.

Here, we introduce a computational framework based on unsupervised AI that replaces verbal descriptors with quantitative, data-driven representations, placing histology on a reproducible quantitative foundation. It is interpretable and intuitive as it transposes to histology the mathematical formalism and methods of RNA sequencing transcriptome analysis.

We demonstrate our framework with the construction of an atlas of human histology. In contrast to the idealized healthy tissues depicted in histology textbooks, this atlas rests on the exhaustive analysis of 23,887 H&E slides from the organs of 946 donors of the GTEx dataset^5,6^. It provides a survey of histology spanning both sexes, an age range of 21 to 71 years and— although GTEx call the donors ‘non-diseased’—a number of sub-clinical conditions.

Our quantitative framework makes it possible to compute associations between the expression of morphemes—the histological operational analogs of genes—and all the other data modalities available in GTEx. These include basic anthropometrics variables, pathology notes, clinical data, genomes, transcriptomes, and for a selection of organs, telomere length measurements, methylomes and proteomes. While some associations identified by our framework are expected or already documented, the vast majority are novel. They form a corpus of annotation of human histology unprecedented in size and scope.

The atlas of human histological diversity is intended as a hypothesis-generating research resource to explore human histological variation. It is delivered as a highly interactive web site, available at <u>histologyatlas.ulb.be</u>, covering 2,560 morphemes from 40 human organs. It is equipped with tools to explore the visual dimensions of morphemes, survey their associations with other data modalities and search them with any molecular, pathology and clinical terms documented in GTEx. It is also provided as flat files empowering users to run their own computational investigations. Most analyses showcased in this article stem from interactive atlas exploration followed by custom computations.

## Results

### A simple framework for quantitative histological expression analysis

Two concepts form the core of our framework: morphemes—the histological computational equivalent of genes—which are categories of small unit tissue area, and reference histological atlases, which are the histological computational analogs of annotated genomes.

To quantify the histological expression in a whole slide image (WSI, Fig 1a), we first cut it into thousands of tiles, our units of tissue area. Second, each tile is embedded by a pretrained AI vision model into a morphological space. In that space, tiles with similar morphologies are close to one another. Third, tiles are assigned to morphemes based on their embedding vector and a reference atlas built beforehand as explained below. Fourth, we count the number of tiles assigned to each morpheme within the WSI of interest.

**Figure 1.**
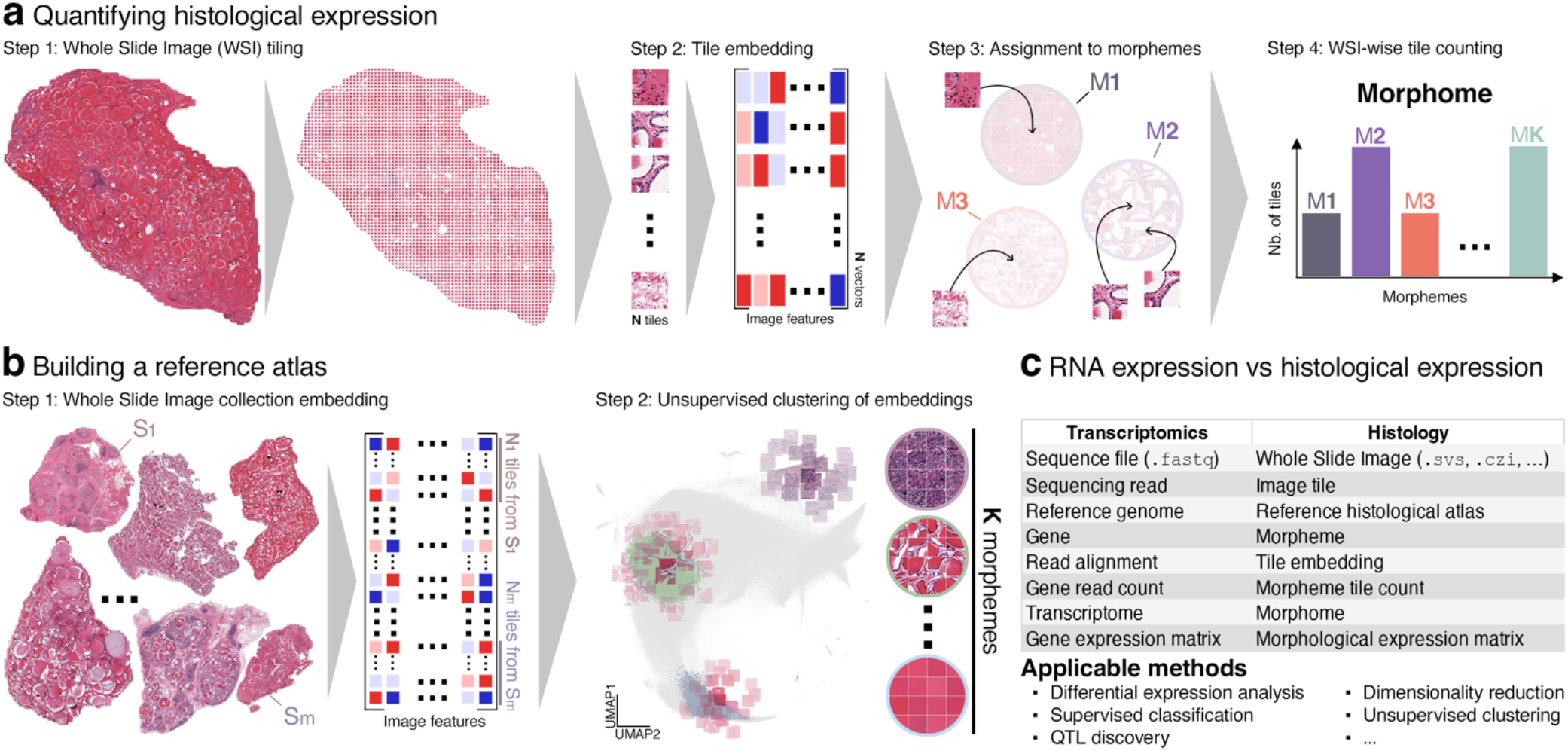
A simple framework for quantitative histological expression analysis. **a**, Quantification of the histological expression of a WSI. **b**, Atlas construction. **c**, Comparison of gene and morpheme expression.

Hence, morpheme expression is defined as an area measurement—convenient units are mm^2^ or % of tissue surface—whose resolution is set by tile size. We call morphome of a WSI the vector of expression values of all its morphemes. Computationally, it is to histology what the transcriptome is to RNA expression.

To build a histological atlas (Fig. 1b), we first apply steps 1 and 2 of the quantification pipeline to a large reference corpus of WSI. We then cluster the embedding vectors with unsupervised clustering. The resulting clusters define the morphemes for this reference corpus that we subsequently use to quantify histological expression in individual WSI (Fig 1a). Because our approach is unsupervised, the slides to be quantified and the atlas must be compatible (see Discussion).

This simple approach to histology quantification follows the bag-of-words principles originally designed for text retreival^7^ and latter applied to images^8^, whereby local descriptors are clustered into a discrete vocabulary and each document/image is represented as a count vector. It presents several advantages: it is automatic, it is not limited by human cognition, and it is interpretable; The core concepts fit operational entities familiar to biomedical researchers: Morphomes and RNA-seq transcriptomes are both count vectors representing the abundance/expression of biological units. A key byproduct is that the analytic framework developed in the last decades for RNA expression can now be applied to histology (Fig. 1c).

### The atlas of human histological diversity

Previous studies have conducted targeted analyses of the GTEx histological slide corpus or a subset of it^9–23^. Based on supervised tissue segmentation in 23 organs, clinical annotations and transcriptomes Cisternino et al.^14^ proposed a more open-ended approach. Here, based on our unsupervised framework we present—and deliver as a web resource—an atlas covering 40 organs and all data modalities available in GTEx.

We generated reference atlases (Fig. 1b) independently for the 40 organs of GTEx for which WSI were available (Fig. 2). Tile size was set to 110×110 μm^2^ and the number of morphemes to 64 per organ (see Discussion). Overall, 23,887 WSI containing 172,742,565 tiles were processed (Fig. 2a). We next measured the morphome of each WSI (Fig. 1a). Hereafter, we refer to morphemes as ‘prostate morpheme M**24**’, for example, to denote the morpheme with ID 24 in the prostate sub-atlas.

**Figure 2.**
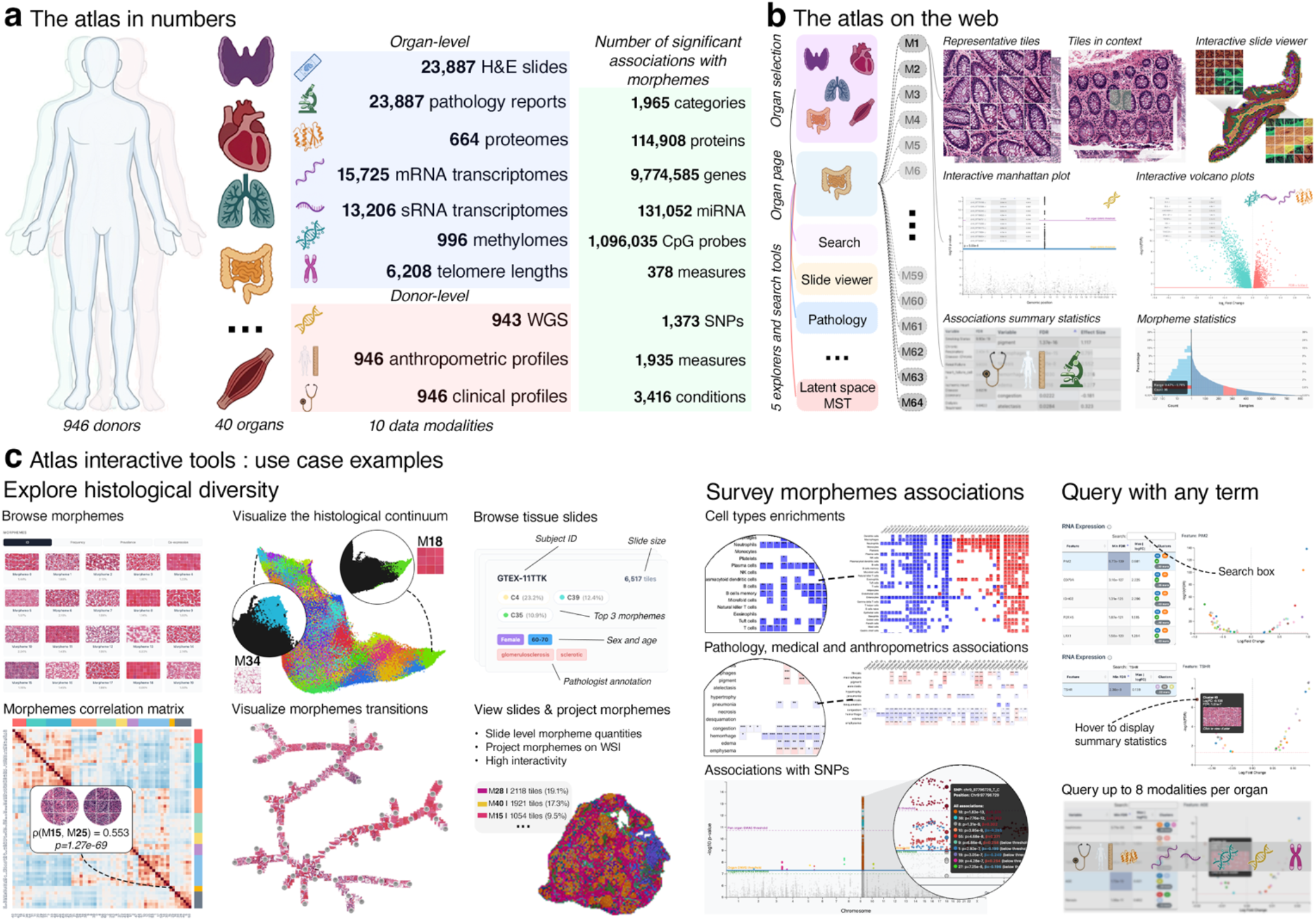
The atlas of human organs histological diversity. **a**, The atlas surveys all organs with WSI available in GTEx. Morphemes were annotated with 11,125,747 significant associations between their expression and 6 sample-level and 3 donor-level data modalities (breakdown by organ in Suppl. Fig. S1, Suppl. Tables S1-S3). Associations were adjusted for age, sex and technical variables. **b**, The atlas is available as an interactive Web site at <u>histologyatlas.ulb.be</u>. It includes 2,560 pages documenting 64 morphemes in 40 organs. They show example morphemes tiles and their surroundings. They include or link to interactive viewers to visualize their position in the histological continuum, their distribution across WSIs and their associations with other data modalities. **c**, The atlas is equipped with interactive tools to explore, survey and query morphemes. Exploration may consist in browsing the morphemes, looking at their co-expression, exploring their relationships in the morphological space via interactive 3D UMAPs or minimum spanning trees. Users may also select slides with specific donors’ characteristics and project morphemes on them with interactive, zoomable, slide viewers. Interactive heatmaps survey association of morphemes with other data modalities per organ. Organ-level search pages power queries of morphemes with terms from any modalities, including gene names.

To annotate the resulting 40×64=2,560 morphemes we computed the associations of the expression of each one with variables from 9 orthogonal datasets available or computable from GTEx (Fig. 2a). These include data modalities matched with individual WSI such as the ‘pathology notes’ recorded by GTEx pathologists and a range of molecular data (Fig. 2a). They also include data modalities matched with individual donors such as anthropometrics, clinical data, genomes and technical confounders (Fig. 2a). The latter are ischemic time, i.e. the time interval separating death from dissection, and the GTEx cohorts (‘organ donor’, ‘post-mortem’ and ‘surgery’). Both have strong associations with histology and other data modalities (see Discussion). Therefore, associations were adjusted for these in addition to age and sex. Overall, 11,125,747 significant associations are reported in the atlas.

The morphemes and their rich annotations were integrated into a Web resource (histologyatlas.ulb.be) that includes tools to interactively explore, survey and search morphemes (Fig. 2b, c), and into downloadable tables for custom investigations.

### The atlas vs. traditional pathology observation

Since the atlas generation is purely unsupervised, its concordance with the categories of pathology is far from granted. We therefore evaluated whether categories extracted from GTEx pathology notes^24,25^ could be predicted with linear modeling from morphemes expression. Among 76 categories eligible for classification (i.e. with >30 cases), 68 (89%) could be predicted with an AUC>0.6, 38 (50%) with an AUC>0.7 and 18 (24%) with an AUC>0.8 (Fig. 3a).

**Figure 3.**
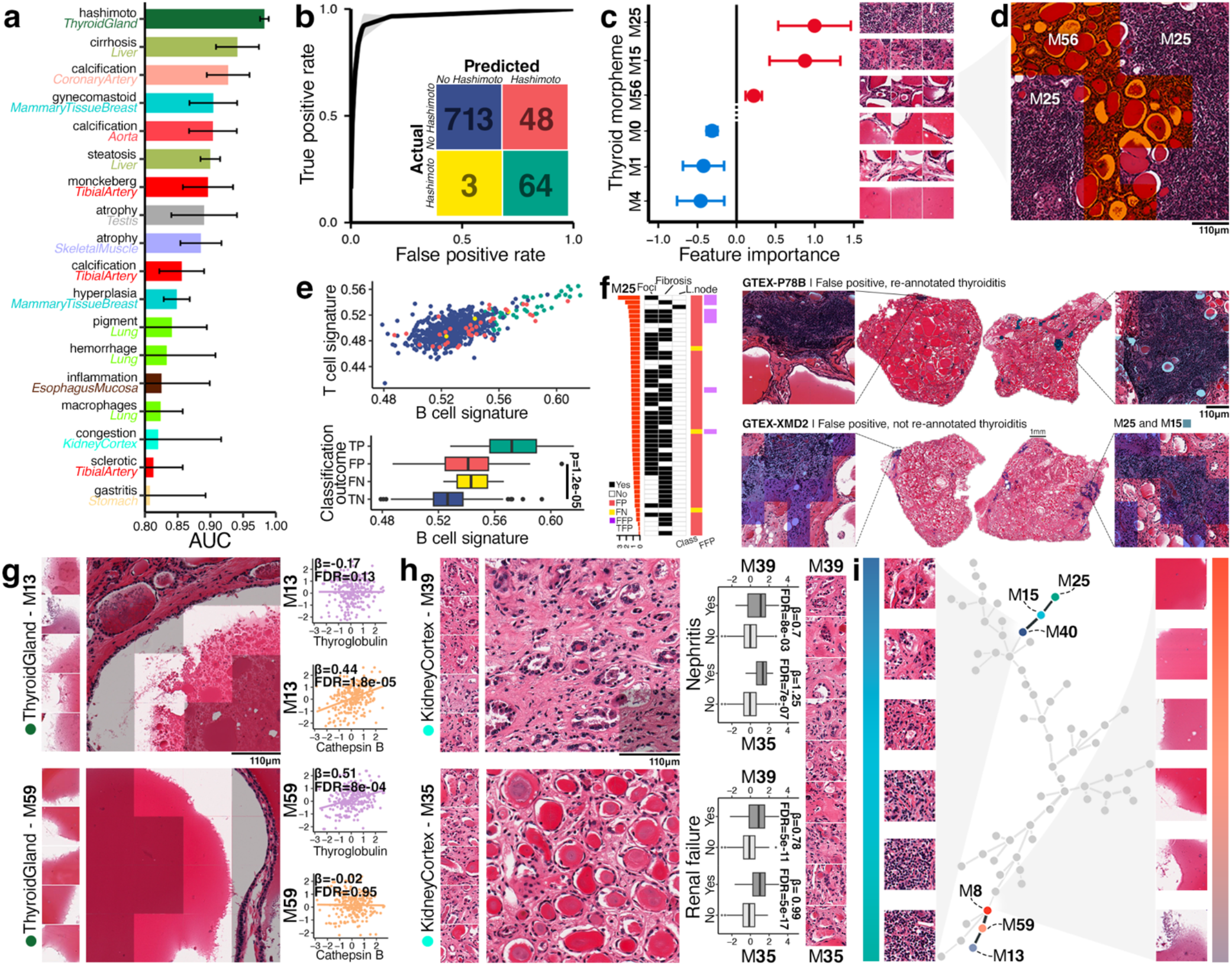
The atlas vs. traditional pathology observation. **a**, Pathology categories predicted with AUC>0.8 (full analysis in Suppl. Table S4). **b**, Performances of the thyroiditis classifier. **c**, Top morphemes in thyroiditis prediction. Three sample tiles are shown for each morpheme. **d,** Tiles from M**56** (shaded in orange), the 3^rd^ most important predictor, cover follicles, not immune foci. **e**, upper panel, Each point represents a WSI with the *x* and *y* axes depicting the expression of B and T cell RNA signatures, respectively, in its associated transcriptome. True positives are enriched in T and, even more so, B cells. Lower panel, distributions of the B cell RNA signature by classification status. **f**, Re-evaluation of lymphocytic thyroiditis diagnostic of misclassified slides (details in Suppl. Table S5). The slides are ranked by decreasing M**25** expression (tiles of M**25** shown in panels c and d). Labels depicted in the 3 black and white columns were assigned during re-evaluation. The orange/yellow column show the original error type, the purple/white column depicts our class re-assignment. Most FP were confirmed as errors (TFP), a few were found to be true positive (FFP). Two slides with comparable M**25** expression are shown: one was reassigned to the thyroiditis group, not the other—illustrating the quantitative inconsistency of classical evaluation. **g**, Thyroid morpheme M**13** and M**59**. For each, representative tiles are shown as a vertical strip next to a wider 4×4 tile context (tiles not in the morpheme shaded in grey). The associated expression of thyroglobulin and cathepsin B with both morphemes are shown on the left. **h**, Kidney cortex morphemes M**39** and M**35** and their association with nephritis and renal failure. The left strip shows tiles along the shortest path in the kidney cortex morphological space linking the centroids of M**39** and M**35**. **i**, Minimum tree spanning the 64 thyroid morphemes. Tiles along the two branches highlight an immune (blue to green) and colloid (red to purple) continuum.

To further understand how morphemes predict pathology categories, we examined in detail the Hashimoto lymphocytic thyroiditis classifier (AUC=0.98±0.0066, Fig. 3b). The top discriminative morphemes (Fig. 3c) were morphemes 25 (M**25**) and M**15**, both containing immune infiltrates, as expected. Interestingly, tiles from M**56**, the third most predictive feature, were centered on follicles. These tiles did not contain obvious immune infiltrates, but their surrounding tiles did (Fig. 3d). Thus, the atlas captured follicular changes associated with lymphocytic thyroiditis.

We next investigated the misclassified cases (Fig. 3b). False positive samples were enriched in T, and even more so B cells RNA signatures, although not as much as true positives (Fig. 3e). We further reviewed misclassified slides (Fig. 3f). Two of the 3 false negatives were relabeled as normal. Among 48 false positive, 6 were reclassified by a board-certified pathologist as *bona fide* thyroiditis and 28 contained dense lymphocytic aggregate, but in numbers insufficient to qualify as thyroiditis (Fig. 3f). Yet, 15 false negative samples expressed more M**25**, i.e. lymphoid foci tiles, than two of the reclassified samples (Fig. 3f). This illustrates how our framework may help improve on qualitative judgement. In addition, neglecting smaller, less numerous, foci is debatable as additional aggregates may be present in the larger 3D volume. Overall, while lymphocytic thyroiditis prevalence was estimated^24^ to 8% of GTEx donors, the morpheme-based quantitative predictions led to a prevalence of dense immune foci of at least 12%. GTEx does not includes anti-TPO antibody measurements. Thus, a clinical parameter that completes the definition of chronic thyroiditis is unavailable to interpret the pathology notes and atlas-based estimates. Cohorts with quantifications of dense lymphocytic morphemes matched with anti-TPO measures are the natural next step to calibrate the quantitative approach against clinical disease.

The atlas captures histological nuances not easily grasped by traditional pathology examination. For example, thyroid M**13** and M**59** both contain tiles with a transition from colloid to empty white space (Fig. 3g). But this transition is sharper in M**59** than in M**13**, which also shows a disordered textures and debris suggesting colloid degradation. Both morphemes are detectable in >98% of thyroid slides, but in the low frequency range (∼1% of tiles), and show very weak co-expression (Pearson’s r=0.1). We asked whether this histological nuance could be explained by protein expression. The expression of M**59**, but not M**13**, increases with that of thyroglobulin (TG, Fig. 3g). Conversely, the expression of morpheme M**13**, but not M**59**, increases with the expression of cathepsin B (CTSB, Fig. 3g). CTSB expression is typical of active TG endocytosis/resorption and intraluminal predigestion in hormone-releasing follicles^26^—in agreement with the visual signs of colloid degradation. Surprisingly, M**13** was also associated with cathepsin Z (CTSZ, Suppl. Fig. S2a), a carboxymonopeptidase expressed in monocytes with no documented activity in thyrocytes. Accordingly, the mRNA cell type signature enrichment analyses available in the atlas point to a mild neutrophil infiltration (Suppl. Fig. S2b), suggesting that M**13** is expressed in a context of mild inflammation. Overall, the atlas reveals subtle histological nuances and provides molecular context to interpret them.

Some morphemes document known morphologies not recorded in GTEx pathology notes. For example, kidney ‘thyroidisation’, a kidney morphology with uromodulin-filled tubules visually similar to thyroid follicles in 2D^27^, is captured by kidney cortex M**35** (Fig. 3h). Morphemes are discrete categories needed for expression measurement, annotation and verbal communication, yet they are built from the essentially continuous morphological space. To visualize the morphological continuum connecting kidney cortex M**35** and another morpheme, M**39**, we displayed tiles along the shortest path in morphological space between the centroids of M**35** and M**39** (Fig. 3h). Both morphemes are associated with pathology categories ‘nephritis’ and diagnostic of renal failure, but associations are 3-8 orders of magnitude stronger for M**35**, suggesting its association with more advanced disease. We hypothesize that the continuum shown Fig. 3h depicts a trajectory from simple tubular atrophy to thyroidisation.

This approach to the histological continuum was systematized by drawing minimal trees spanning morphemes across the morphological space of each organ. The web atlas provides interactive viewers to examine these trees and project annotations onto them. As an illustration, Fig. 3i shows the minimal tree spanning the thyroid morphological space. Some branches highlights, for example, distinct gradients of colloid textures, or immune cell density (Fig. 3i).

In summary, the atlas complements existing approaches to histology by providing quantitative accuracy, extensive molecular context and a handle on the morphological continuum.

### Histological expression is associated with anthropometrics and lifestyle

The atlas confirmed the major impact of age on histology reported in earlier studies of GTEx slides^18,28–30^ (Fig. 4a), but unveiled a more detailed picture because it includes orders of magnitude more histological categories. In addition, the extensive morpheme annotations paint more accurately the complex anthropometric and clinical contexts associated with aging-related morphemes. For example, the two morphemes most positively associated with age in the aorta were M**18**, expressed in the *tunica media* and *adventitia,* and M**0**, expressed in the *tunica intima* (Suppl. Fig. S3). M**0** was more strongly correlated with ischemic heart disease, calcification, osteopondin expression and macrophages infiltration (Suppl. Fig. S3). These associations are consistent with the presence of cholesterol crystals in M**0** tiles (Suppl. Fig. S3). This suggests, that M**18** reflects a healthier aging path than M**0**.

**Figure 4.**
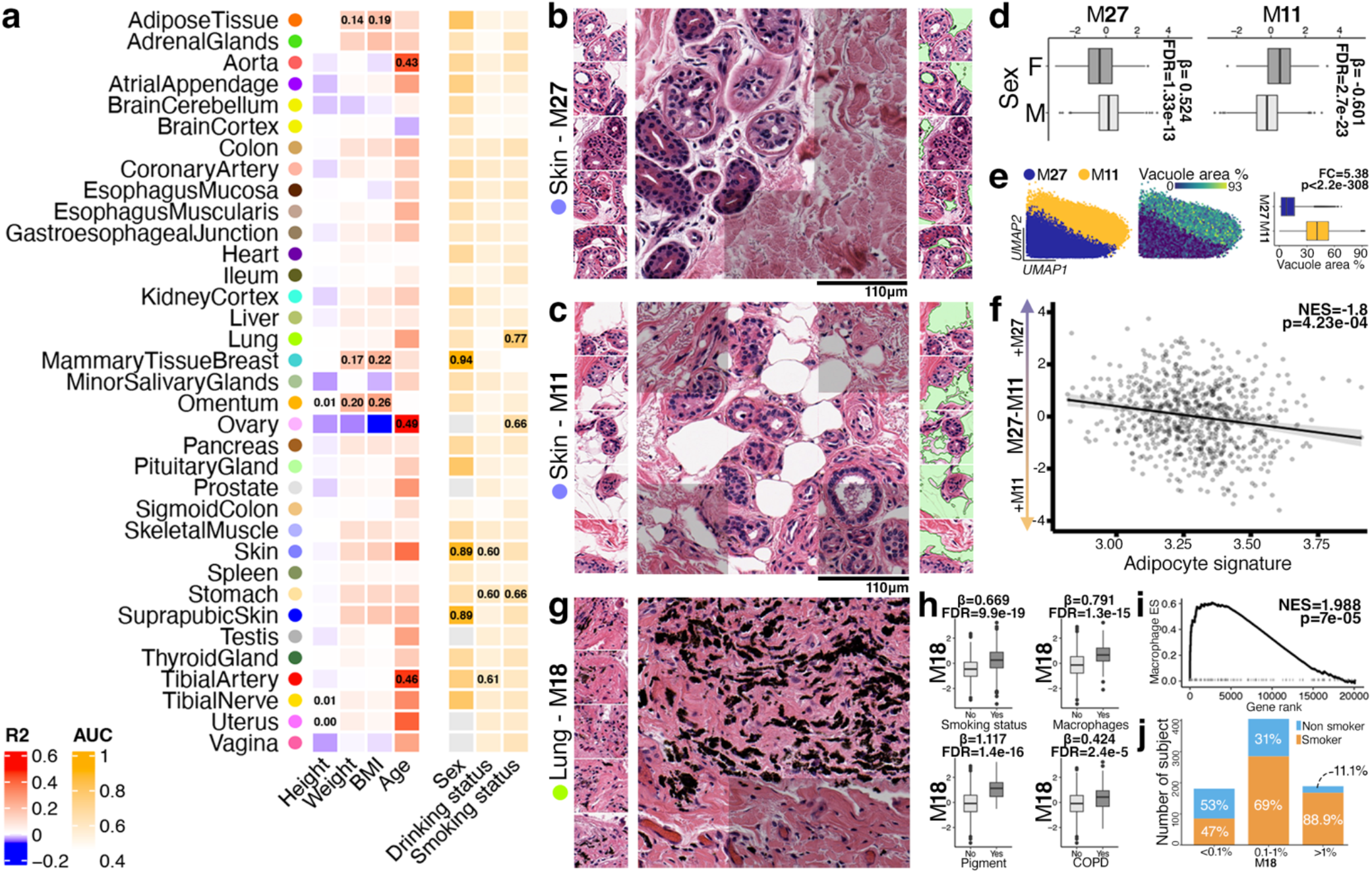
Histological expression is associated with anthropometrics and lifestyle. **a**, Prediction of donors’ height, weight, BMI, age, sex, drinking and smoking status from morphemes (Suppl. Table S4). Top 3 AUC (discrete variables) or R^2^ (continuous variables) are highlighted for each variable. **b**, Left, 5 tiles from sun-exposed skin morpheme 27; Center, 3×3 tile context of a M**27** tile (M**27** tiles not shaded); Right, same 5 tiles as left, but with adipose vacuole segmentation in green. **c**, same as panel b for M**11**. **d**, M**27** is more expressed in males, M**11** in female. Data were adjusted for age. **e**, Left, zoom in the M**27**/M**11** boundary in the sun exposed skin morphological space: it is very sharp; Center, same colored by adipose vacuole area; Right, distribution of adipose vesicle areas in tiles from M**27** and M**11**. **f**, Each point depicts a sun-exposed skin WSI-matched transcriptome. The more M**27** is expressed, the less enriched an adipocyte RNA signature is. The enrichment calculation included adjustment for age and BMI, ruling out their effects. **g**, Five representative tiles and one 3×3 tile context of lung Morpheme M**18**. **h**, Left, M**18** is more expressed in smokers and COPD donors. It is associated with pathology reports of pigment and macrophages. **i**, The latter is confirmed by the over-expression of macrophage signature. **j**, The proportion of smokers increases with M**18** expression, but it is present at sizable levels in non-smoker.

Sex predictions reveal the pervasive presence of moderate sex-related signals (predicted with AUC>0.6 in 27 organs and AUC>0.7 in 8 organs, Fig. 4a). Breast was the most predictive organ followed by skin, subcutaneous adipose tissues and the pituitary gland (Fig. 4a). Specific examples are morphemes M**11** and M**27**, the best predictors of female and male sexes, respectively, in sun-exposed skin. They both contain eccrine sweat glands (Fig. 4b, c). M**11** is overexpressed in females, M**27** in males (Fig. 4d). As visual inspection suggested more adipocytes around sweat glands in M**11**, we quantified adipocytes vacuole area: it is 5.4-fold higher in M**11** (Fig. 4e). Furthermore, the expression difference M**27**–M**11** is negatively associated with a mRNA adipocytes signature enrichment adjusted for BMI (Fig. 4f). Finally, projecting individual tiles’ lipid vacuole area onto the sun-exposed skin morphological space revealed a crisp boundary fitting tightly the M**11**/M**27** boundary (Fig. 4e). Thus, these morphemes represent eccrine glands with sharply distinct mechanical and paracrine microenvironments, yet their differential expression between females and males shows a strong bias, but not a binary difference (Fig. 4d).

The best predictions for sex/age-adjusted BMI were obtained from omentum (R^2^=0.26±0.05), breast (R^2^=0.22±0.04), subcutaneous adipose tissue (R^2^=0.19±0.04), adrenal glands (R^2^=0.18±0.02) and suprapubic skin (R^2^=0.16±0.04) morphemes. Weight was less predictable than BMI (best R^2^=0.2±0.04), height was not predictable (best R^2^=0.005±0.01). BMI associations with subclinical conditions are addressed in Fig. 6.

We next investigated the predictability of two life-style variables, smoking and drinking, from histology. Surprisingly, morphemes from skin and tibial artery had predictive power for drinking status comparable to that of stomach and higher than those of other segments of the GI tract and liver. In contrast, lung dominated smoking status predictions, reflecting the direct effect of tobacco on this organ. For example, lung morpheme M**18** exhibited massive associations with GTEx donors’ smoking status, and with pathologist’s reports of pigments and macrophage infiltration (Fig. 4h). Anthracosis is indeed readily visible in M**18** tiles and M**18** expression is associated with an enrichment of the macrophage RNA signature (Fig. 4i). The proportion of smokers increased with M**18** expression (Fig. 4j). Remarkably, among donors with M**18** expression >0.1%, 25% were labelled as non-smokers (Fig. 4j). The survey and search tools of the atlas revealed many more systemic effects of tobacco. An example is the negative association of smoking status with transverse colon M**47** expression, a morpheme characterized by dense immune patches (Suppl. Fig. S3). This finding is consistent with the decreased rate of ulcerative colitis in smokers^31^. Another example is aorta morpheme M**50** (Suppl. Fig. S3). It shares many features of M**0** discussed earlier, including association with heart disease, calcification and macrophage infiltration, but in contrast to M**0**, it is expressed in the *tunica media* (Suppl. Fig. S3). In addition, M**0** associates most strongly with age, M**50** with smoking status (Suppl. Fig. S3), suggesting different driving mechanisms.

Thus, our quantitative framework reveals novel aspects of histological aging, sexual dimorphism and the histological impact of drinking and smoking.

### Genomic variations associated with histological expression

We conducted a Genome-Wide Association Study (GWAS) to detect histological quantitative trait loci (hQTL), i.e. loci associated with morpheme expression. We considered three distinct significance thresholds accommodating different atlas use cases focused on a single morpheme, all morphemes of an organ, or the entire atlas (see Discussion). This yielded 287/6/2 morphemes associations with 1,105/80/35 unique SNPs distributed among 267/3/1 genomic peaks at genome-wide/organ-wide/atlas-wide significance (Fig. 5a, Supplementary Tab. S6). Two examples are showcased below. Other hits can be explored interactively in the online atlas.

**Figure 5.**
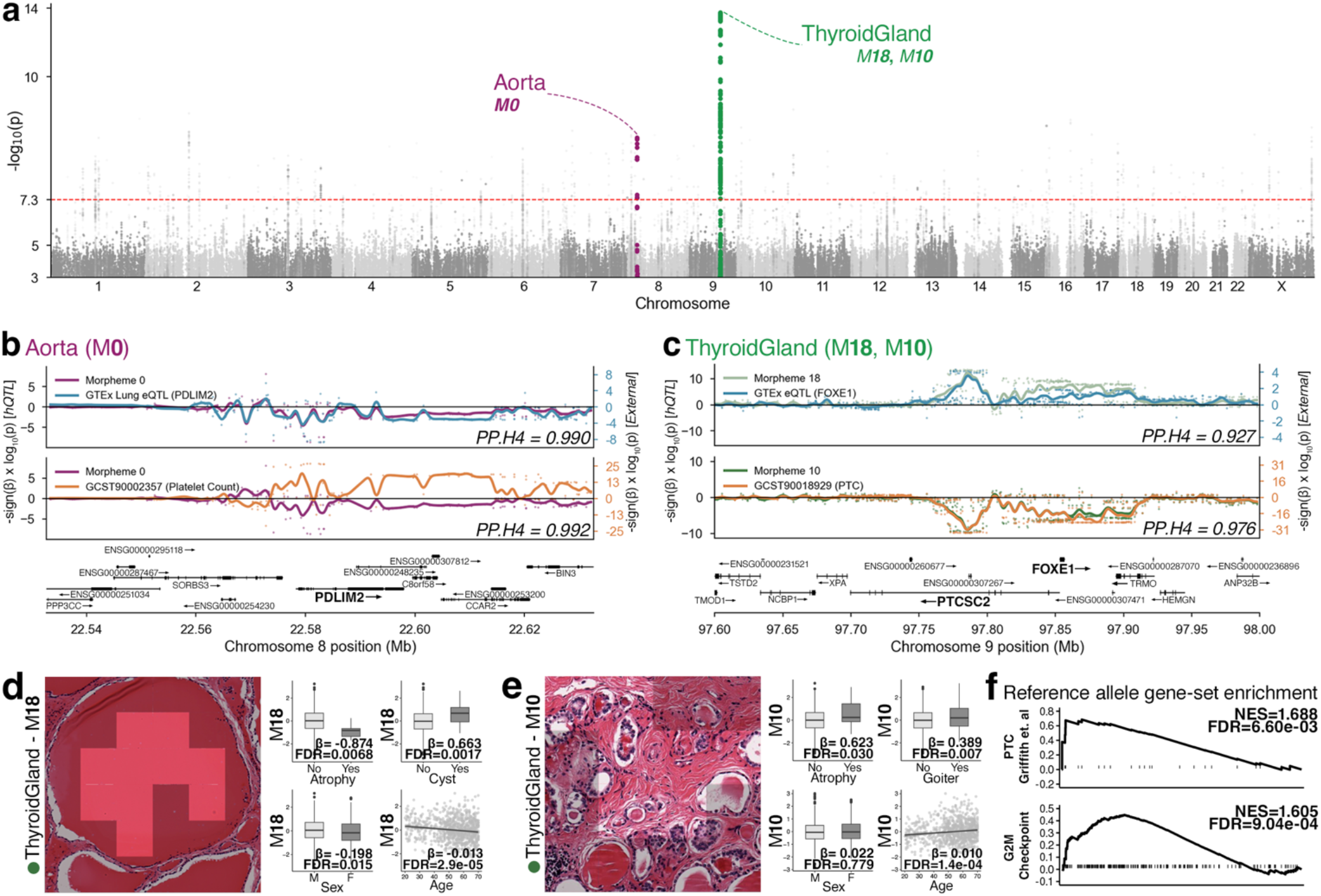
Histological quantitative trait loci (hQTL). **a**, Pan-organ Manhattan plot for hQTLs (Suppl. Fig S6). **b**, Aorta morpheme M**0** is associated with SNPs overlapping PDLIM2. The signal colocalizes with that of PDLIM2 eQTLs in the lung, a site of platelets production, and with signal from a previous GWAS^58^ (GWAS Catalog ID: GCST90002357) covering blood platelet counts. PP.H4 measures the probability that traits are associated and share a single causal variant. PP.H4>0.8 is considered strong evidence. **c**, Two thyroid morphemes, M**18** and M**10** associated with SNP overlapping *PTCSC2* and *FOXE1*. The signal colocalizes with thyroid eQTL for *FOXE1* and a previous GWAS on PTC risk^37^ (GWAS Catalog ID: GCST90018929). The sign of the associations is identical for M**10** and PTC risk, which are opposite to the M**18** and the *FOXE1* eQTL signals. **d**, M**18** tiles (non M**18** tiles greyed out) contain pure colloid. This, morpheme is positively associated with pathology diagnostic of cyst and negatively with atrophy and tends to be up-regulated in males and younger individuals. **e**, In contrast, M**10** is associated with aging, goiter and atrophy. Associations with age, sex and pathology categories are presented Suppl. Fig S4 for all lead variant-associated morphemes. **f**, A RNA signature composed of genes over-expressed in papillary thyroid cancer (PTC) and a G2M checkpoint signature are significantly enriched with the M**10**-associated reference allele.

The aorta morpheme M**0**, described earlier in the context of aging and calcification, is associated with a locus overlapping PDZ and LIM domain 2 (*PDLIM2*; Fig. 5b). Association of *PDLIM2* with platelet count is documented in the GWAS catalog and this signal significantly colocalized with ours (Fig. 5b). Our signal did not colocalize with the GTEx *PDLIM2* eQTL signal in the aorta, but it did in the lung (Fig. 5b), a major site of platelet production^32^. The bone marrow could not be investigated as it is not part of GTEx. Platelets are mediators of inflammation and arterial calcification^33^. Overall, the atlas documents at least two types of aorta calcification: M**50** driven by smoking in the *tunica media* and *adventitia*, and M**0** driven by aging and genetics in the *tunica intima* (Suppl. Fig. S3).

The most significant GWAS hit across the atlas (Fig. 5a) linked several thyroid morphemes with a region on chromosome 9 that included Papillary thyroid cancer susceptibility candidate 2 (*PTCSC2*) and Forkhead box E1 (*FOXE1*) (Fig. 5c). *FOXE1* is a major transcription factor in thyroid development^34^. *PTCSC2* is a lincRNA antisense of *FOXE1*^35^, and a modulator of its expression^36^. Our GWAS signal colocalized with the thyroid *FOXE1* eQTL signal and the signal of a previous PTC-related GWAS^37^. The same locus was associated with large follicles— interpreted as suggestive of goiter—in a previous study^38^, but the finer granularity of the atlas and its annotation led to a significance two orders of magnitude stronger and enriched the interpretation. Six morphemes associated positively with the alternative allele contained tiles with pure colloid (Fig. 5c), in agreement with the large follicles reported in ref. 38, but they tended to be positively associated with diagnostic of cyst and negatively associated with goiter, atrophy, age and female sex (Fig. 5d and Suppl. Fig. S4). In contrast, two morphemes associated with the reference allele and were positively correlated with goiter, and with aging (Fig. 5e and Suppl. Fig. S4). Moreover, RNA signatures of PTC^39^ and proliferation were over-expressed with the reference allele (Fig. 5f). Taken together, these results suggest that big follicles associated with the alternative allele—which is also the major allele—are phenotypes antagonistic to cancer. The minor allele, on the other hand, is associated with morphemes characteristic of goiter and fibrosis, and suggestive of a precancerous state. Two recent genetic studies also reported a lower cancer risk associated to the alternative allele^40,41^, but found it to be associated with higher goiter risk. We conjecture that this apparent contradiction may arise from the differences in presentation between clinically detectable and occult postmortem goiter in GTEx slides.

### System histology: pan-organ histological groups

To reveal structures of morpheme expression across organs, we constructed for each morpheme of each organ predictors from the morphemes of other organs. A clustering based on organ pairs morpheme-wise mean R^2^ is presented Fig. 6a. Subparts of organs clustered together, for example, transverse and sigmoid colon, sun-exposed and suprapubic skin, so did the major segments of the GI tract.

**Figure 6.**
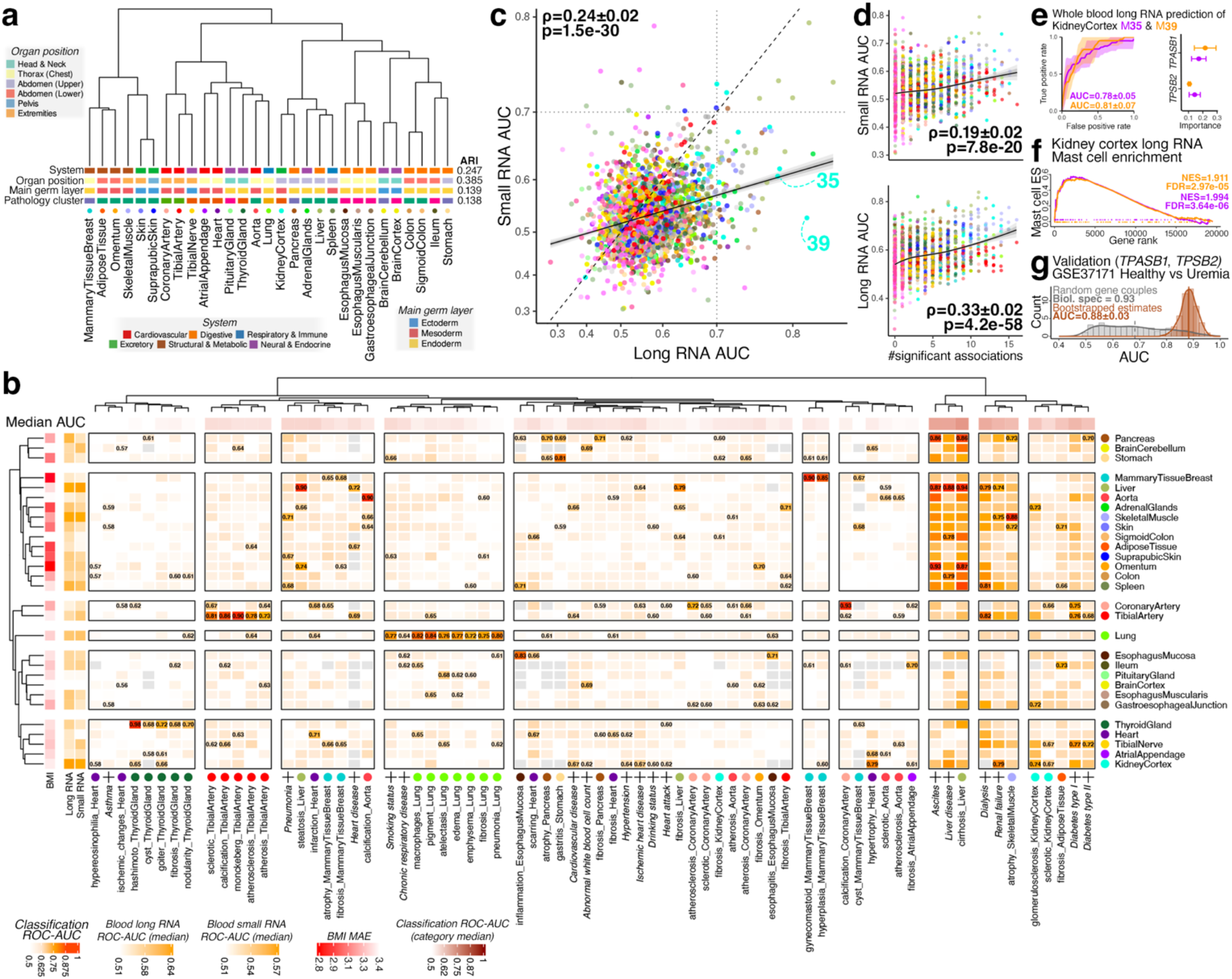
System histology. **a**, Clustering based on cross-organ predictability of morphemes. The Adjusted Rand Index (ARI) measures the concordance between the clustering and the organ classification systems depicted as horizontal tracks. It is bounded in [0, 1], 1 means perfect concordance. **b**, Predictability of BMI, pathology categories and clinical variables (labeled ‘+’) from the morphemes of different organs (Suppl. Table S4). The median of the AUC across organs (top track) is higher for conditions with systemic effects. The long RNA and small RNA tracks depict the median AUC of organs’ morphemes prediction from blood transcriptomes. **c**, Long vs small RNA for morpheme predictions from blood transcriptomes (Suppl. Fig. S5 and Suppl. Table S7). Kidney cortex M**35** and M**39** are highlighted. **d**, The AUC increase with the number of associations of morpheme with diseases-related pathology categories. **e**, Feature importance and AUC of *TPSB2* and *TPSAB1* in Kidney cortex M**35** and M**39** predictions. **f**, Enrichment of a mast cell RNA signature in the kidney is associated with the expression of M**35** and M**39**. **g**, Validation of *TPSB2* and *TPSAB1* as markers of uremia in an independent data set, GSE37171^56^. The brown histogram depicts the distribution of 1,000 bootstrapped AUCs. The grey one stands for the distribution of AUCs for 1,000 randomly selected sets of two genes. This controls for the biological specificity of the prediction (Suppl. Methods and ref. ^59^): owning to the systemic effect of kidney diseases (panel b), many genes predict uremia, i.e. the grey distribution is spreads over high AUCs, but *TPSB2* and *TPSAB1* predict uremia better than random genes. Specificity is 0.93. This metric varies between 0 (not specific) and 1 (all random genes yield AUCs inferiors to *TPSB2*/*TPSAB1* AUCs).

To further characterize the factors driving cross-organ histological correlations, we compared this clustering with three organ classification systems reflecting embryology, function, and anatomy (Fig. 6a). Morpheme-based clustering was weakly concordant with the dominant germ layers, slightly more with the organs systems, and moderately concordant with body location (Fig. 6a). Beside these systemic organizing factors, diverse specific effects may contribute to the picture. For example, the clustering of tibial artery and nerve likely result from the presence of shared connective tissues in imperfectly dissected blocks, and the proximity of thyroid and pituitary is consistent with the TSH/T3-T4 feedback loop coupling these two glands.

Having examined the general cross-organ morpheme expression trends, we next focused on cross-organ predictions of organ-specific pathology categories and donor-level clinical terms (Fig. 6b). First, the organs clustering based on these predictions weakly matched the clustering based on morpheme predictions (Fig. 6a), suggesting that the signals shown Fig. 6b are not primarily driven by dominant germ layer, organ system and body location. Second, pathology categories of an organ were generally better predicted by morphemes of that organ. Surprisingly, however, 39% of cross-organ predictions were better or on par with intra-organ predictions. For examples, heart hypertrophy and ischemic heart disease were better predicted by kidney cortex than by heart or cardiovascular system morphemes, adipose tissue fibrosis was better predicted by ileum and skin morphemes.

Cross-organ predictions analysis quantified and delineated the body-wide impact of major metabolic syndromes. Liver disease, cirrhosis and ascites were predicted with AUC>0.70 by morphemes in up to 21 organs spanning all organ systems. This profile was related to that of dialysis/renal failure and muscular atrophy which were, in addition, also predictable from tibial nerve and artery. Diabetes type I and II clustered together and with glomerulosclerotic kidney and fibrotic adipose tissues. Nuances were visible, with type I most predictable from tibial nerve, tibial and coronary arteries, and type II from Kidney, tibial nerve and pancreas. On a broader scale all these liver/kidney/diabetes-related conditions clustered together, and the associated morphemes included the best sex and age-adjusted BMI predictors (Fig. 6b).

We next asked whether morpheme expression could be predicted from blood long and short RNA transcriptomes, i.e. clinically accessible material. Seventy-two morphemes could be predicted with AUC>0.7 from long RNA, 19 from short RNA, 8 from both (Figs. 6b and c). AUCs were higher for morphemes significantly associated with pathology and clinical disease terms (Fig. 6d). Most morphemes predicted with AUC>0.7 belonged predominantly to the kidney and the liver, and were associated with related diseases (Fig 6c), in agreement with the systemic impact of disfunctions in these organs (Fig 6b). For example, kidney cortex morphemes M**35** (thyroidisation, Fig. 3h) and M**39** (diabetic and glomerulosclerotic, Fig. 3h) were predicted from long RNA with balanced AUC=0.81±0.07 and 0.78±0.05 (Fig. 6e), respectively. The two best features were tryptase beta-2 (*TPSB2*) and tryptase alpha/beta-1 (*TPSAB1*), two mast cell-specific proteases. M**35** and M**39** were among the three kidney cortex morphemes most enriched for the mast cell RNA signature (Fig. 6f). Thus, the mast cell signal in the blood mirrored their infiltration within the kidney cortex. Mast cells association with chronic kidney disease (CKD) is well documented^42^. While serum tryptase protein level is mostly used as a marker of anaphylaxis, other mast cell disorders^43^, and hematologic diseases^44^, it has received limited attention in the context of CKD^45^. The TPSB2 and TPSAB1 gene combination predicts uremia in an independent dataset^46^ and the predictive signal is biologically specific (Fig. 6g).

Overall, our quantitative framework enables previously intractable organism-wide histological analysis.

### Technical discussion

The atlas rests on data and procedures that come with their own biases, limitations and distortions. Understanding them is critical to properly interpret our atlas and to develop future atlases. Here we discuss some technical aspects of GTEx and atlas construction and highlight open research questions. Several of the points we raise also apply to previous studies based on GTEx histology slides^9–23^ (Suppl. Table 9) and, in several instances, gene expression.

While sub-clinical conditions are already a pervasive theme in the atlas, their prevalence is likely underestimated: GTEx recruitment excluded subjects with BMI<18.5 or >35 (according to ref. 25, but we found 11 donors with BMI>35); surgical protocols favored macroscopically normal tissue; pathology notes reporting is very limited in some organs (Supp. Table S2); slides represent only tiny fractions of organs’ 3D volume, thus small focal lesions are missed with high probability; and overtly cancerous slides were excluded^25^. In addition, diseases representation is likely biased by an over-representation of older Caucasian males.

GTEx samples were collected postmortem, with post-mortem interval (PMI) <24hrs. PMI was predictable from morphemes with R^2^ up to 0.59 ± 0.06 (Suppl. Fig. S6a), a result relevant to forensic. Previous PMI predictions from RNA-seq were slightly better^47^, but H&E slides are more tractable and interpretable. Protein expression was also measurably affected, which may help understand tissue dissociation. For example, thyroid M**63** is positively associated with PMI, M**57** negatively. Their member tiles overlap average-sized follicles, but epithelial sheets from M**63** were less cohesive and its junction proteins content was decreased compared to M**57** (Suppl. Fig. S6b). Organ donors were typically younger. Associations with PMI and donor type are documented and were adjusted for (Suppl. Fig. S6e). Yet, residual confounding effects of PMI and related variables (Suppl. Fig. S6c) may persist. Restricting the atlas to transplant donors would have further dampened these effects, but it would also have eliminated aging-related signals. For example, the *PTCSC2* signals presented in the previous section vanishes in GWAS restricted to transplant donors (Suppl. Fig. S6d). All statistical associations we presented were adjusted for ischemic time and cohort, and when appropriate age and sex. Importantly, addressing zero-inflation (Supplementary Methods) is a prerequisite for effective ischemic time and cohort adjustment (Suppl. Fig. S6e).

Block dissection often included non-parenchymal tissues, potentially biasing associations. Attempts to remove them, however, did not improve GWAS signals. Moreover, their distinction from pathological features can be ambiguous. For example, adipocytes in a thyroid slide may result from thyroid lipomatosis (Suppl. Fig. S7a) or inaccurate dissection (Suppl. Fig. S7b).

Slide tiling, which is part of most histology IA pipelines, may produce a continuum in the embedding space. An example is the tiles covering the otherwise crisp dermis/epidermis transition forming a continuum of dermis-to-epidermis ratio in the skin morphological space (Suppl. Fig. S8). The minimum spanning tree representations available in the web atlas helps sorting apart tiling-induced from biologically relevant continuums.

Tile size controls the magnification at which the atlas surveys histology. We kept it fixed for simplicity, but no single magnification optimally reveals all aspects of histology. For example, we built our atlas from tiles of 110×110 μm^2^, but a tiling at 220×220 μm^2^ results in a thyroid *PTCSC2* GWAS signal 4 orders of magnitude larger, because larger tiles more accurately capture signal related with large colloid patches and therefore large follicles (Suppl. Fig. S9). By contrast, another morpheme had PTCSC2 associations much less dependent on magnification (Suppl. Fig. S9). The informational dependencies across different magnifications and how to integrate them while preserving interpretability are open problems.

The AI tile embedding model defines the morphological space. Morphemes’ definitions depend on the coarse structure of that space. We deployed many vision models over the years developing the atlas and found it not to be a critical parameter. Models trained/fine-tuned on WSIs have finer histological discriminatory power which leads to better performances in supervised linear probing benchmarks^48^. But we observed that a significant fraction of models pretrained on common internet images were texture discriminators able to distinguish tiles from different organs and producing thyroid morphemes capturing most major structures within organs. We compared the thyroid morphemes derived from three embedding models (Suppl. Fig. S10): DINO ViT-B/8^49^ a 86 million parameters model trained on ImageNet^50^, a collection of common internet images; Lunit-DINO^51^ and Virchow2^52^, fine-tuned with self-supervision protocols on histopathology images. Lunit-DINO is a 22 million parameters model trained on 15,672 WSI from TCGA (the model selected to produce the atlas). Virchow2 is a 632 million parameters model trained on 3.1 million WSI^52^. The morpheme correlation structure is reproducible, overall, across models (Suppl. Fig. S10).

Visual variation induced by slice thickness, staining, scanning and other variations in WSI acquisition collectively generate batch effects which impact the representations produced by vision models. In the supervised setting, the classifiers added on top of the embedding model may learn batch-independent classification rules for the target classification problem. But batch effects are an unsolved problem in the unsupervised setting. This has two implications. First, our atlas is possible because the unified GTEx standard operating procedures (SOP) produced a slide collection with minimal technical variation^25^—although isolated cases of staining variations are noticeable in the atlas, for example Brain Cortex M**26** includes tiles from a single WSI with a unique hue. In contrast, technical variation dominated over biological variation when we attempted to build an atlas from TCGA THCA (thyroid cancer) WSI, because it is multicentric and not unified by SOPs (Suppl. Fig. S11). As a result, morphemes were specific of production centers. Pathologists see biologically relevant features despite major technical variation. Endowing vision models with this ability is a major open challenge.

Morphemes are anchors partitioning the morphological space. They are shaped by the clustering algorithm and the granularity, *K*, at which we discretize the morphological space. In the absence of clear information-based cut-off (see elbow plot Suppl. Fig. S12), and because we did not tailor the atlas for a specific research question, we set *K* to 64, a fine granularity compared to previous unsupervised approach to histology with GTEx^9,56,28,38,29,18^. Importantly, this setting grants atlas users the option to create coarser morphemes by summing existing ones, while maintaining a tractable atlas size. *K*=64 does incur some over-clustering. For example, brain cortex morphemes M**42** matches a faint and biologically meaningless focus nuance (Suppl. Fig. S13). On the other hand, many subtle variations revealed at *K*=64 are molecularly and/or clinically grounded as exemplified with thyroid morphemes M**13** and M**59** and aorta’s M**0** and M**50** (see previous sections). Beyond histology, we contend that mathematical innovations are needed to accurately handle and interpret biological continuums without resorting to any form of discretization.

Morphemes were annotated with bulk molecular profiles. It is nevertheless informative and often consistent with their local spatial expression, for example the top morphemes associated with insulin RNA in the pancreas is Langerhans islet morpheme M**15**. Yet, many RNA/protein-morpheme associations reflect slide-level co-expression, not local effects. While ideally suited to annotate an atlas, omics spatial data are currently not available at GTEx scale.

The atlas is an open-ended resource that should accommodate different use cases. Some users will be interested in specific morphemes, others in all morphemes from an organ or in the entire atlas. We applied Benjamini-Hochberg corrections accounting for proteomes, transcriptomes and methylomes sizes, and for number patho-clinical variable tested. Thus, the atlas will be conservative for users conducting morpheme-level investigations, and not conservative enough for those surveying the entire atlas. In the first situation, the strong signals we observe, in particular for transcriptome associations with typically several thousand of genes associated with a single morpheme, mitigate the conservative corrections. Users in the second situation may further correct the p-values provided as downloadable tables for the number of organs they investigate. We took a slightly different stance in the presentation of GWAS results, since users may run colocalization studies on peaks slightly below genome-wide significance: the significance thresholds are adjustable in the web interface to fit users’ need. Thresholds for genome-wide, organ-wide and atlas-wide significance are displayed as guides.

## Conclusion

We presented a principled approach to histology quantification, applied it to the construction of an atlas of human histological variation with rich multimodal annotations and demonstrated its use as an open-ended hypothesis generating resource.

Our approach is part of a wider epistemological trend in the production of scientific atlases which has been analyzed by Daston and Gallison^57^. Early atlases represented ideal specimens free of natural irregularities. A layer of cognitive bias was later eliminated by mechanical reproducibility, for example photography. Yet, ‘trained judgment’ was still required to sort apart relevant from irrelevant material and produce categories^57^. Our framework aims to reduce this source of human bias in histology. While GTEx scientists had to set inclusion criteria, we processed with equal attention all tissue sections and tiles. Morphemes do rest on technical decisions, but they were derived automatically based on unsupervised and importantly, reproducible, machine learning. Our quantitative framework also enabled automatic annotation of morphemes with layers of information that help ground their interpretation.

The atlas brings nuances to the concepts of normal histology by revealing the diversity associated with variations of age, sex and BMI across organs. Moreover, while many features of GTEx design likely resulted in a significant underrepresentation of samples with subclinical pathologies (see Discussion), their associations with morphemes are surprisingly pervasive in the atlas. The atlas provides a handle to investigate the continuum between health and disease from a phenotypic layer, histology, in which disease states may be visible before they produce clinically detectable symptoms.

Our findings suggest that systemic histological variations related to liver and kidney diseases as well as diabetes, and to a lesser extent other diseases, may be quantifiable from minimally invasive blood tests. This is a step towards specific preventive follow-up and/or treatment, but of course, the prognostic value of these variations will need to be assessed in future studies to address over-treatment. We also demonstrate the existence of measurably different aging paths in the aorta. This highlights the potential of histology to distinguish ‘healthy’ from ‘pathological’ aging trajectories in future aging research.

Finally, we cannot stress enough the importance of research on unselected donors—an ideal approached by GTEx and reached by the UKBiobank and comparable programs—to our understanding of human diversity and its biomedical consequences.

## Data availability

The atlas was built on data available from the GTEx v10 dataset (https://www.gtexportal.org). Specific analyses also used the GWAS catalog (https://www.ebi.ac.uk/gwas/home, studies GCST90002357 and GCST90018929) and GEO (https://www.ncbi.nlm.nih.gov/geo, study GSE37171). The interactive web atlas and underlying tables are available at https://histologyatlas.ulb.be. Temporary reviewer access to morphomes: https://mycloud.ulb.be/index.php/s/dyMDHNsWAXqxCic.

## Code availability

Temporary reviewer link: https://mycloud.ulb.be/index.php/s/dyMDHNsWAXqxCic. As technical details of the atlas web interface are being optimized, this part of the code will be made available later—this part of the code does *not* affect the analyses presented in the paper.

## Acknowledgments

This work is dedicated to the memory of Jacques E. Dumont who empowered us with his enthusiasm for fundamental questions. The atlas would not exist without the anonymous donors who made their bodies available to science, their families, the GTEx consortium and researchers publishing high quality open-source tools. We are deeply grateful to all of them. We thank the High-Performance Computing team from ULB Département Informatique for their constant support. We thank ULB researchers who crash-tested early prototypes of the atlas and shared their expert take on it. We thank Benjamin Beck, Michel Georges, Isabelle Migeotte, Marc Parmentier and Gilbert Vassart for their comments on the manuscript. Michel Georges also advised us on technical aspects of GWAS.

## Authors contributions

Z.Z. conducted the AI-related analyses, morpheme generation and contributed to genetic analysis.

Z.Z. and V.D. generated the initial version of the atlas. H.M. developed the multimodal analyses, the Web atlas, suggested some and performed all targeted analyses and contributed to the genetic analyses. D.S. performed genetic analyses. L.L. reviewed predicted thyroiditis slides and provided histopathology expertise. M.T. contributed to concepts underlying the study. V.D. conceived the study and collected funding, developed early prototypes and wrote the paper.

## Competing interests

The authors have no competing interest to declare.

Supplementary figures and methods

## 1 Supplementary Figures

**Suppl. Fig. S1.**
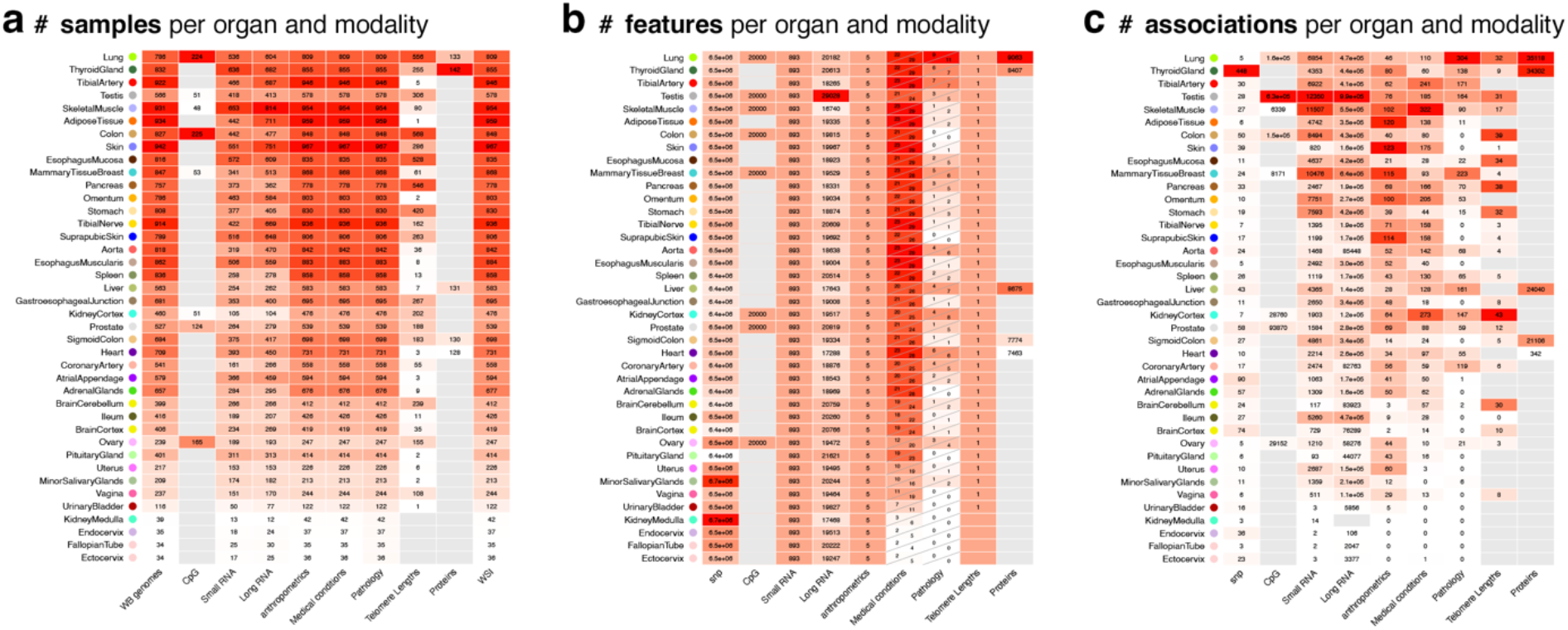
Data, feature and annotation matrices. Colors depict the number of items normalized per column. **a**, Number of WSI for which a data modality (column) is available, per data modality, per organ(rows). **b**, Number of features per data modality, per organ. In columns ‘Medical conditions’ and ‘Pathology’, the top left-side numbers reflect the threshold of at least 15 cases used in the correlation-based atlas annotations. The bottom right-side numbers reflect the threshold of at least 30 cases used in the classification tasks presented in the paper. **c**, Number of significant associations with morphemes per data modality, per organ. Details are available in Supp. Tables S1-S3.

**Suppl. Fig. S2.**
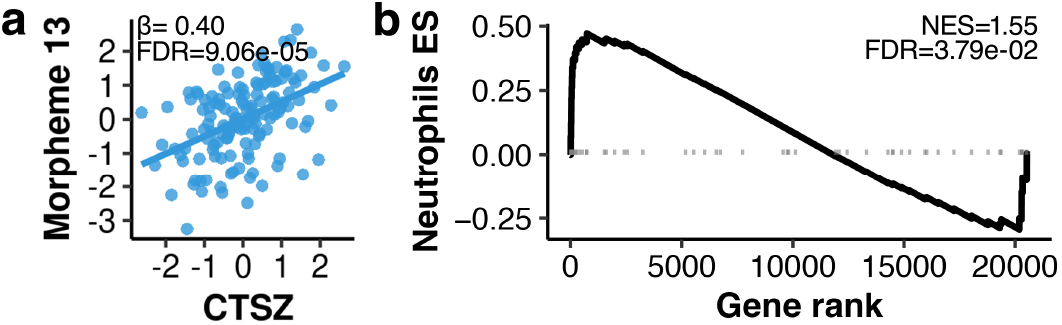
Thyroid morpheme M13 associates with immure-related CTSZ. **a**, CTSZ protein expression is associated with thyroid M**13** expression. **b**, M**13** expression is also associated with the expression of a neutrophil RNA expression signature.

**Suppl. Fig. S3.**
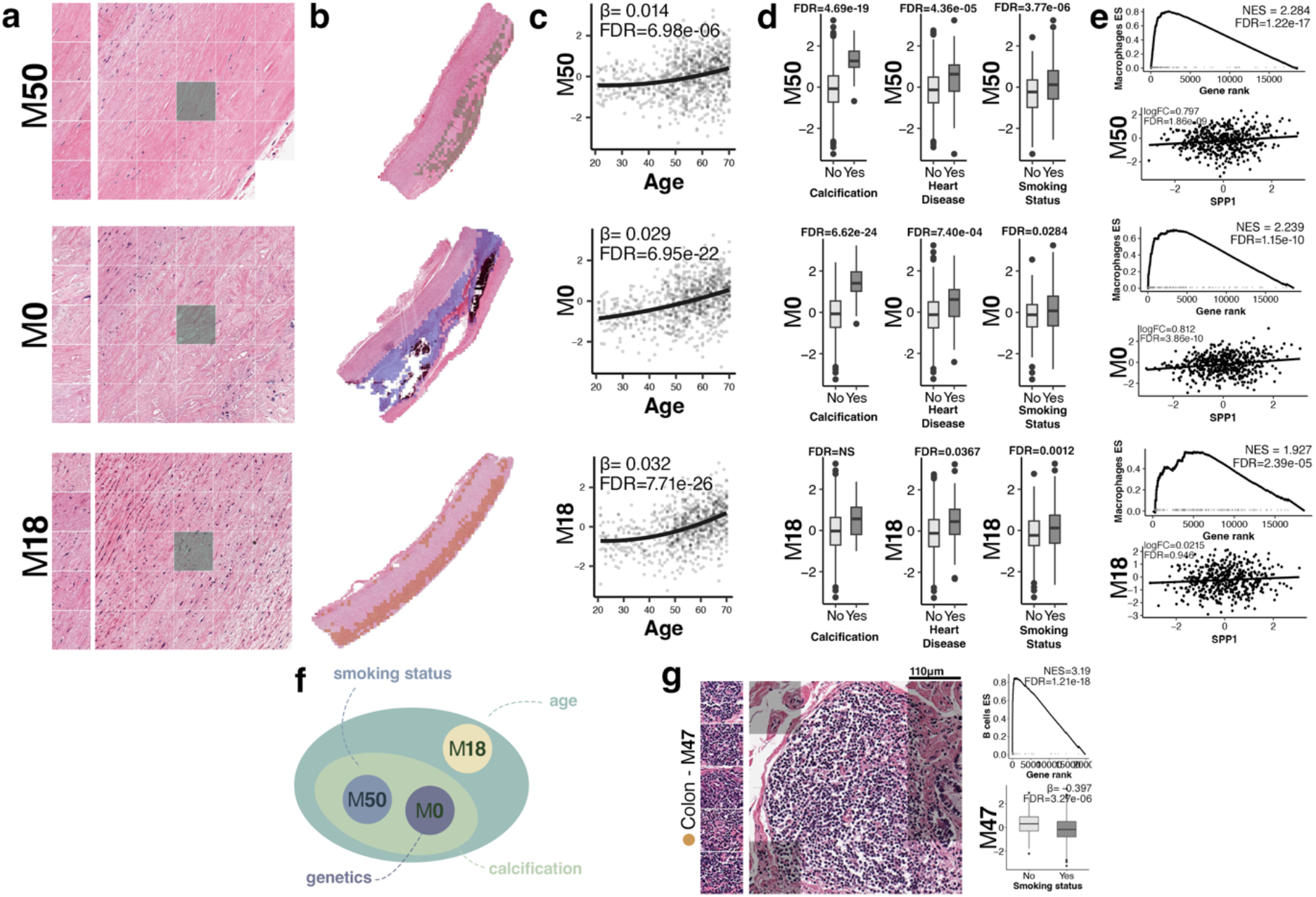
Three aorta and one colon morphemes associated with age and/or smoking. **a**, Five representative tiles and one tile context for aorta morphemes M**50**, M**0** and M**18**. **b**, Same morphemes projected on whole slices. M**50** and M**18** are expressed in the *tunica media* and *adventitia*, M**0** in the *tunica intima*. **c**, The three morphemes are significantly associated with age to different extents, with M**50** being more weakly associated than M**0** and M**18**. **d**, Associations with pathology diagnostic of calcification, and clinical diagnostic of heart disease, and with smoking. M**50** is more strongly associated with smoking. M**50** and M**0** are the most associated with disease. **e**, The enrichment of an RNA macrophages signature is associated with M**50** and M**0** expression. Calcification in these morphemes is confirmed by the up regulation of osteopondin RNA expression (a.k.a. *SPP1*). **f**, Summary diagram: all three morphemes are associated with age, but M**50** less so. M**50** and M**0** are both associated with calcification, but M**50** is driven by smoking, while M**0** is driven by genetics (Fig. 4 in main text). This suggests that M**18** represents a healthy ageing path. **g**, Colon morpheme M**47** is less expressed in smokers and positively associated with a B cell RNA signature.

**Suppl. Fig. S4.**
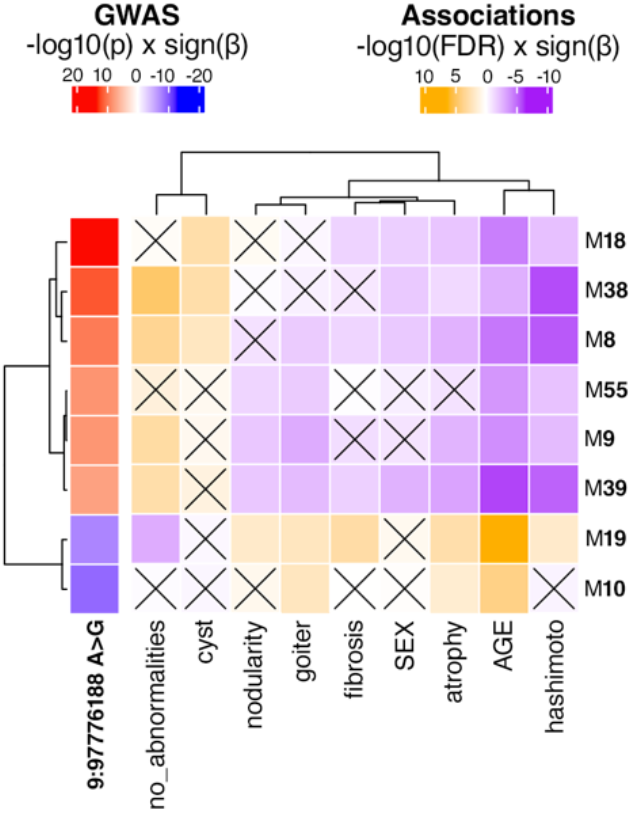
*PTCSC2*/*FOXE1*-associated morphemes vs. age, sex and pathology categories. The first column shows the signed p-value of association with the lead variant on chromosome 9. Crosses denote non-significant associations at *α* = 0.05. Colloid/alternative allele-related morphemes tend to be positively associated with categories ‘cyst’ and ‘no_abnormality’ and to be more expressed in younger donors and females (GTEx encodes males with ‘1’ and females with ‘2’). They also tended to be negatively association with ‘goiter’, ‘atrophy’, ‘fibrosis’ and ‘hashimoto’. A reverse pattern is observed for the two morphemes overexpressed with the reference alleles, M**18** and M**10**.

**Suppl. Fig. S5.**
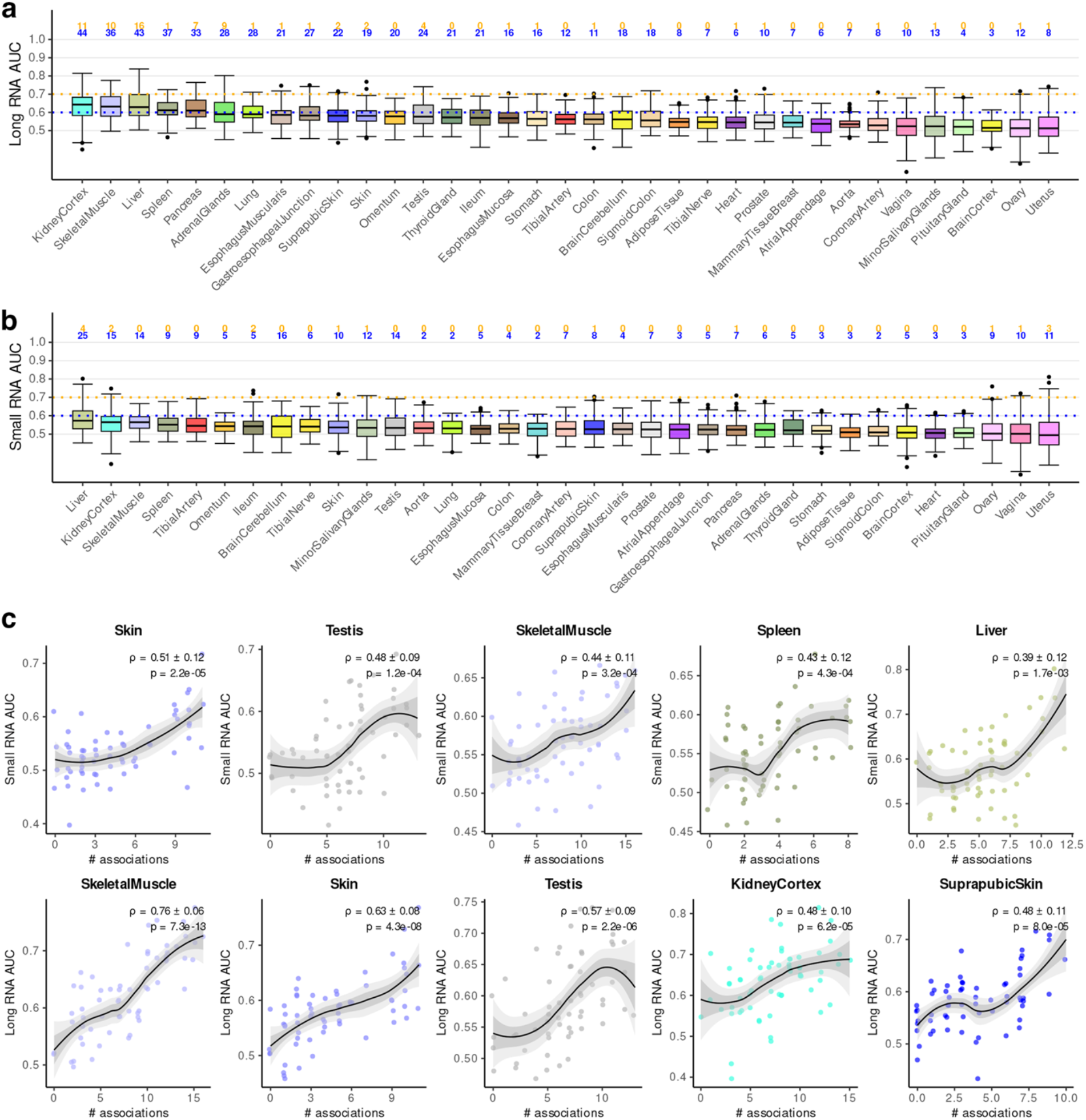
Morpheme predictions from blood RNA. **a**, AUCs for predictions of morphemes from whole blood long RNA. Number of predicted morphemes with AUC>0.6 and AUC>0.7 are provided above each box. **b**, same for small RNA. **c**, Same as main text Fig. 6d for the five organs with highest correlations between number of associations and small and long whole blood RNA-derived AUCs.

**Suppl. Fig. S6.**
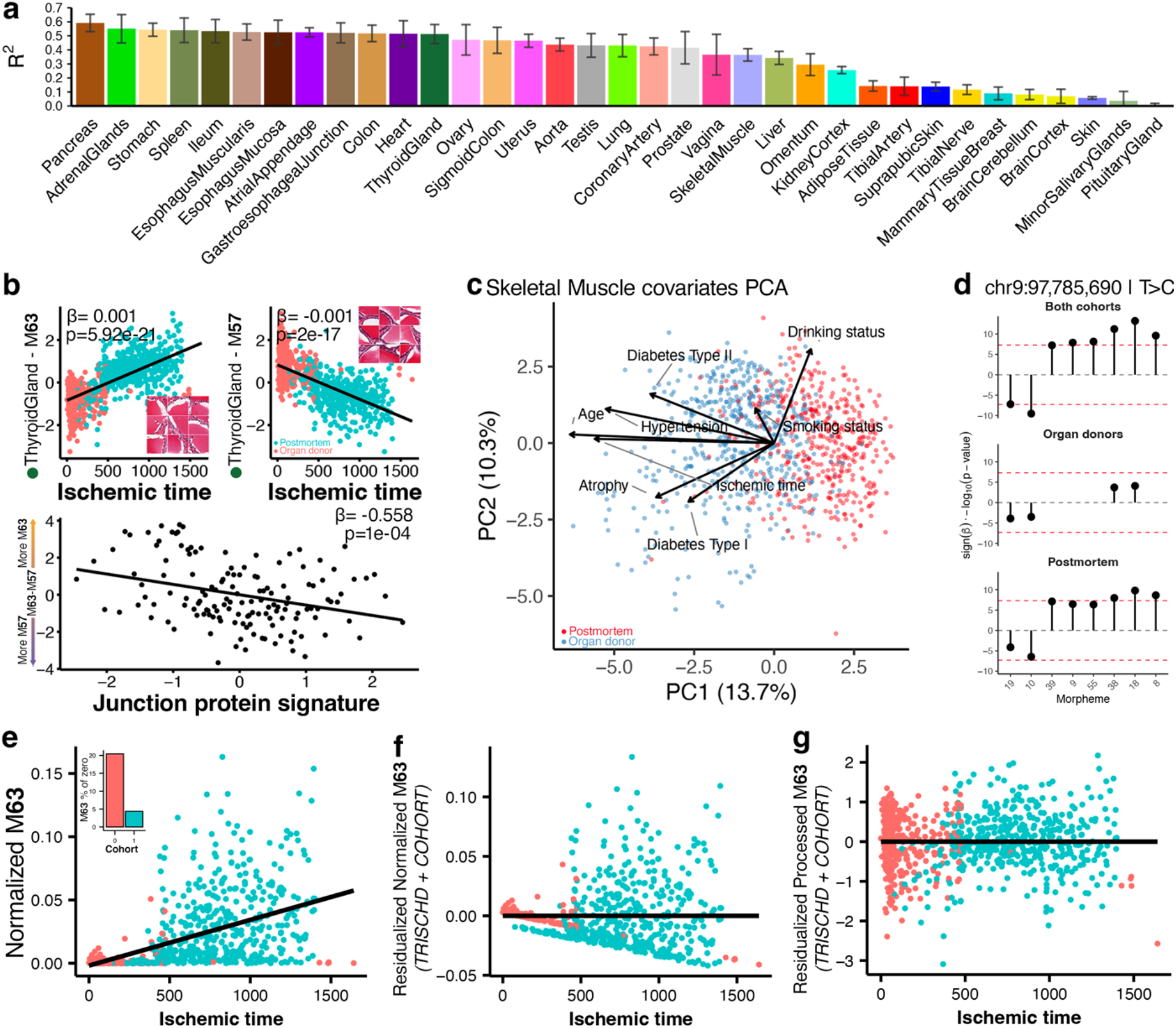
Ischemia is a potential confounder of histology and other data modalities. **a**, Prediction of ischemic time from morphemes. **b**, Tissue dissociation and junction proteins. Top, Ischemic time vs. thyroid M**63** and M**57**. M**63**, but not M**57**, shows loose epithelial sheets. Bottom, protein expression of cell-cell junction genes decreases with M**63** and increase with M**57**. **c**, PCA computed from technical and clinical annotations (labeled on the plot) from skeletal muscle WSIs. Ischemic time confounds the effect of age and age-related conditions. **d**, Association of the lead variant at the *PTCSC2* locus with different thyroid morphemes on the basis the whole dataset or the main cohorts, ‘organ donors’ and ‘postmortem’. The signal is much weaker in in GWAS restricted to the typically younger organ donors. **e**, Normalized M**63** expression shows a positive correlation with ischemic time, but interpretation is confounded by differential zero-inflation between cohorts (inset: ∼20% zeros in ‘organ donors’ vs ∼5% in ‘post mortem’, same color code as in panel c) and unequal ischemic time distributions. **f**, Residuals from a linear model (M**63** ∼ TRISCHD + COHORT) remove the overall trend but retain heteroscedasticity and zero-inflation artefacts, indicating inadequate correction on normalized counts. **g**, After multiplicative replacement and inverse normal transformation, residuals show stable variance across ischemic time. This preprocessing removes zero-inflation artefacts while preserving compositional constraints.

**Suppl. Fig. S7.**
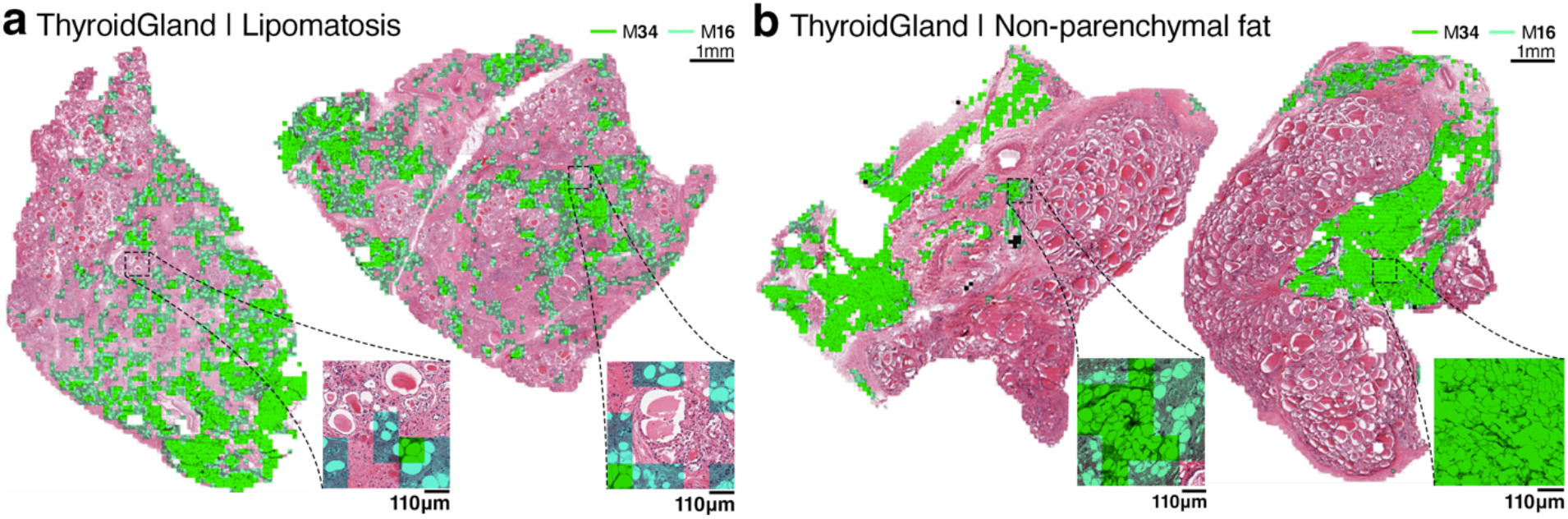
Adipocyte inside and outside the thyroid. Thyroid morphemes M**34** and M**16** capture adipocytes. We projected them on two slides (green shading). **a**, In some slides they are distributed in the parenchyma in a pattern typical of thyroid lipomatosis. **b**, In other slides they are found in residual adipose/connective tissues resected with the organ. These two expression patterns illustrate that removing M**34** and M**16** would eliminate unwanted tissues but also strike lipomatosis out of the atlas.

**Suppl. Fig. S8.**
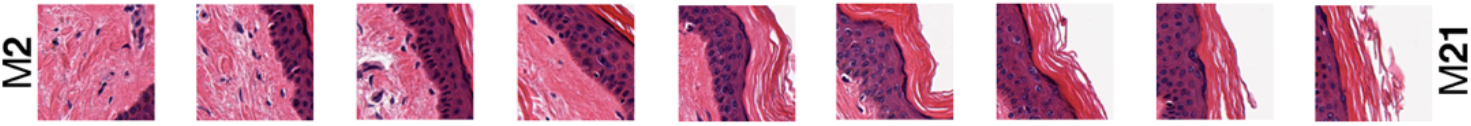
Tiling may induce the illusion of a continuum. This path across the sun-exposed skin morphological goes from dermis (M**2**) to epidermis and outer space (M**21**). The proportions of these three entities within tile vary continuously along that path. Thus, the transition between them is sharp, but tiling creates the illusion of a continuum.

**Suppl. Fig. S9.**
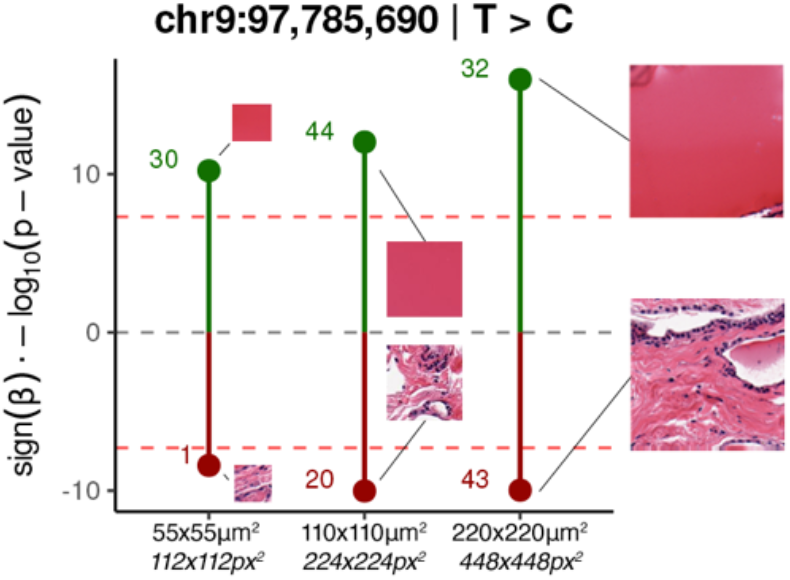
The impact of tile size is context-dependent. Association of *PTCSC2* lead variants with morphemes defined a tile size of 112×112 pixels^2^ (55×55 μm^2^), 224×224 pixels^2^ (110×110 μm^2^) and 448×448 pixels^2^ (220×220 μm^2^). The strength of association increases more dramatically with the colloid morpheme because big colloid tiles are better proxies to capture follicle size.

**Suppl. Fig. S10.**
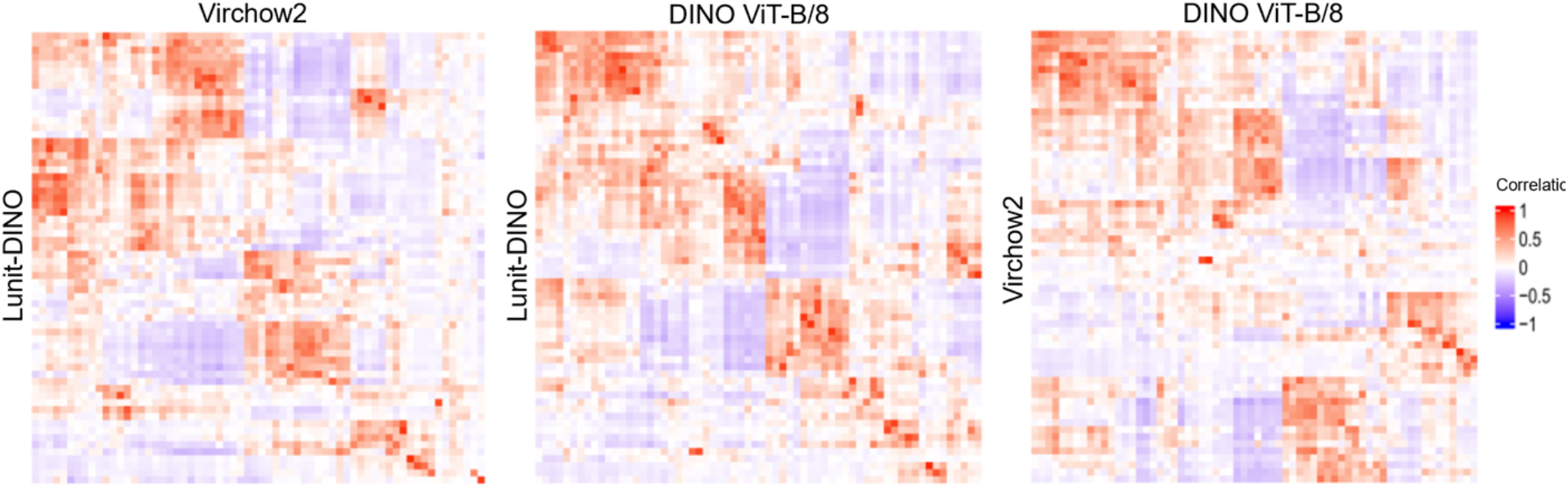
Embedding model convergence. The heatmaps show the correlation between the expression of morphemes generated from clustering with *k*=64 from three embedding models. While not perfectly concordant, some consistent structures are present, matching our qualitative observation that models capture convergent representations of morphology. For example, colloid, immune foci, fibrosis and small degenerate follicles are captured by the three models. Since Gaussian Mixture Model clustering could not be executed on our servers (limited to 1.5TB of RAM) for the 7,100,194 thyroid tiles in the 2,560 dimensions of the Virchow2 embedding space, we used instead the 100 first PCA components of each model.

**Suppl. Fig. S11.**
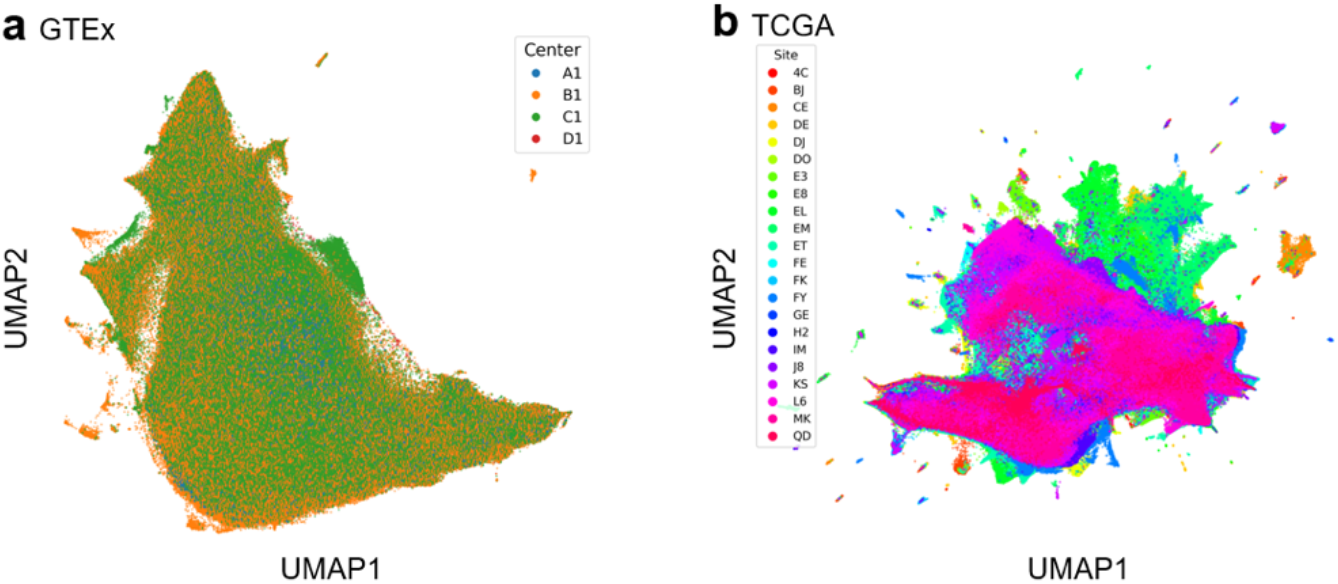
Batch effects in the unsupervised settings. UMAPs computed from tile embeddings and colored by production centers. **a**, GTEx thyroid tiles. Tiles from WSI associated with different centers are intermixed. **b**, TCGA papillary thyroid cancer dataset^1^ (a.k.a. THCA). Tiles tend to group by center. Ten percents of the tiles are shown in each UMAP (i.e. ∼700k for GTEx, ∼3M for TCGA/THCA).

**Suppl. Fig. S12.**
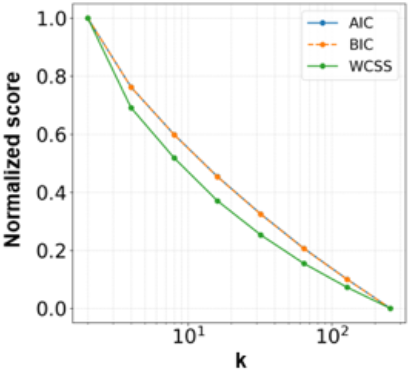
No clear information-based cut-off determines the optimal number of morphemes. This scree plot for the Gaussian Mixture Model clustering of thyroid tiles demonstrates that there is no ‘elbow’—a classical set point for the number of clusters—in the curve. An ‘elbow’ may appear at a much higher granularity. But, WSI-specific morphemes were found at K=128, and even more so at K=256 (not shown) and would only become more numerous as K increases. Morpheme present in one or in very few slides offer little to no prospect of annotation from statistical analysis.

**Suppl. Fig. S13.**
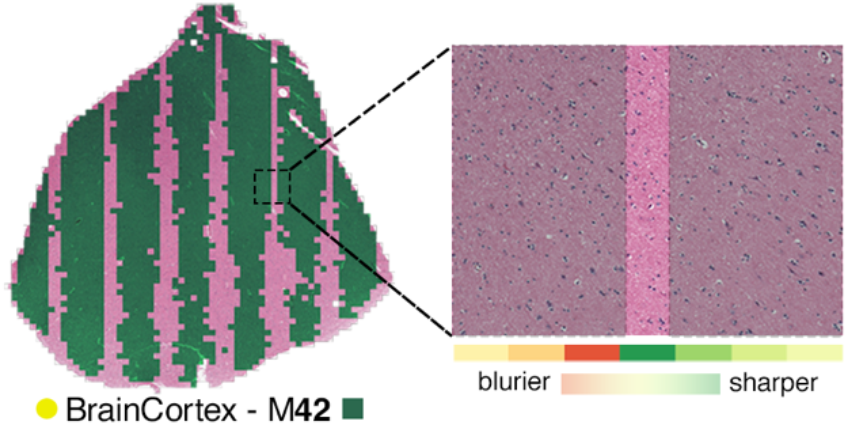
An example of overclustering. Brain cortex morpheme M**42** forms stripes on the slide. These match areas blurred by poor focus. The bottom bar shows the degree of blurring. Specifically, to assess focus quality across whole slide image tiles, we computed a frequency-based sharpness score using Fast Fourier Transform (FFT) on local image blocks defined as the ratio of high-frequency to total spectral energy. We then excluded the sharpest regions (top 20th percentiles, typically including nuclei boundaries) assumed to represent tissue and computed the median sharpness of the remaining background pixels for each tile to evaluate overall focus quality. The full resolution slide with M**42** projection is available here: https://histologyatlas.ulb.be/viewer/BrainCortex/GTEX-11DXY/index.html.

## 2 Methods

### 2.1 Whole slide image selection

All available whole slide images (WSI) were downloaded from the GTEx Globus server. The GTEx project used two methods for tissue preservation: PAXgene fixation (25,288 slides) and dry ice preservation (152 slides). Each histological slide was evaluated by the GTEx quality control team for donor eligibility, ensuring inclusion of only healthy, cancer-free individuals and exclusion of samples exhibiting substantial autolysis. The majority (23,961 slides) were classified as ‘Acceptable’. To ensure dataset uniformity and quality, slides labeled as ‘Unacceptable’ and those processed with the dry ice fixation method were excluded. Additionally, one slide with trichrome staining was removed (GTEX-P44H-1226).

### 2.2 Tissue segmentation

**Fig. M1.**
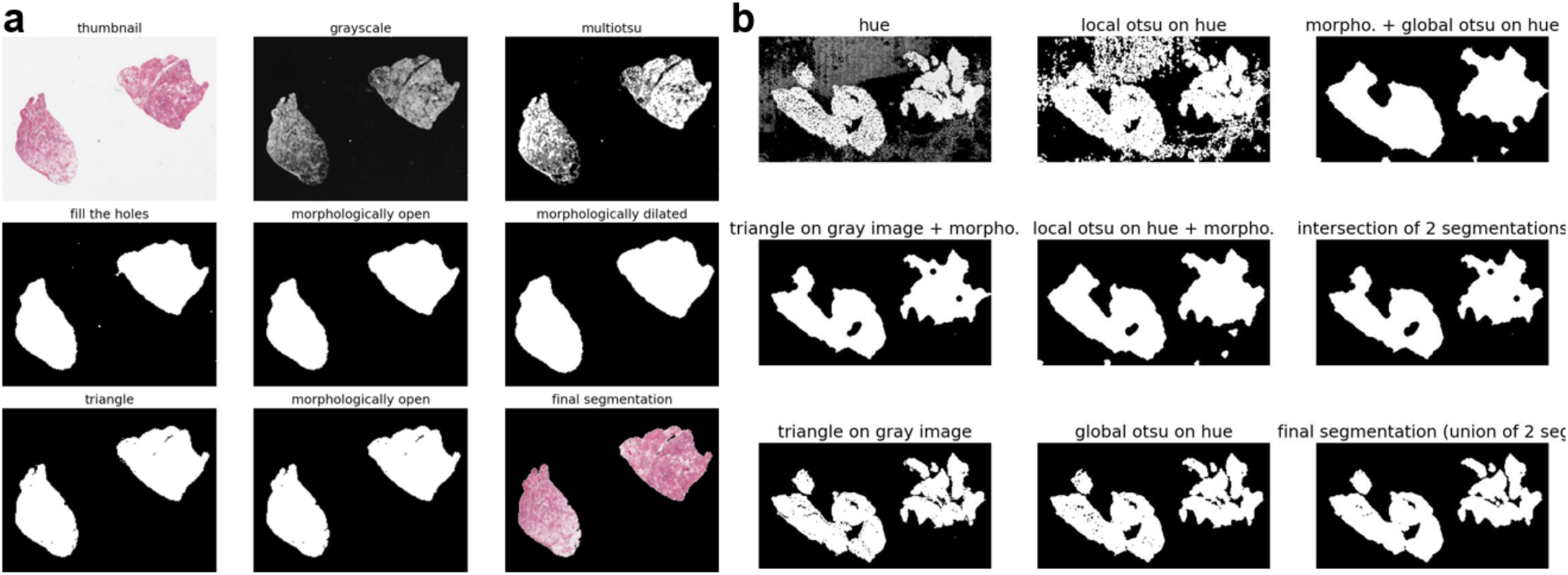
Tissue segmentation. **a,** Segmentation pipeline for non-fatty tissues, in which tissue regions are readily distinguishable from the white background in grayscale space and segmented using Otsu-based thresholding followed by further refinements. **b**, Segmentation pipeline for fatty tissues, where low tissue contrast in grayscale necessitated segmentation in color space; local Otsu thresholding was applied to the hue channel of HSV-converted images, followed by further refinements.

Segmentation of whole-slide images (WSIs) involves an inherent trade-off between (i) preserving all tissue, including low-intensity regions such as lipid-rich (fatty) areas, and (ii) removing preparation- or scanning-related artefacts. Global grayscale thresholding (e.g., Otsu) is generally effective for retaining non-fatty tissue while suppressing many small artefacts, but it can remove genuine tissue when tissue intensity approaches background levels, a frequent issue for fatty regions. Conversely, softer thresholding approaches (e.g., Triangle) better preserve faint tissue but tend to retain background artefacts. Under the working assumption that most artefacts are located in the background, we implemented a two-stage workflow for all slides. Stage 1 performed an Otsu-based coarse tissue–background separation, estimated background statistics, and produces as its direct output a synthetic-background reconstructed image (background replaced by a uniform value). Stage 2 applies softer, artefact-tolerant thresholding on the reconstructed image to refine the final tissue mask. We applied the same segmentation strategy across slides, while parameter choices were adapted to image content as described below. Our segmentation pipeline is based on `scikit-image` version 0.19.3.

*Non-fatty tissues: grayscale-based segmentation (Fig. M1a)*

*Stage 1*. For slides in which tissue intensity was clearly separable from the white background (typically non-fatty samples), coarse tissue detection was performed on low-resolution grayscale images using multi-level Otsu thresholding (`filters.threshold_multiotsu`). Multi-level Otsu was preferred over binary Otsu because the additional class increased sensitivity to low-intensity tissue regions.

The initial tissue mask was refined using sequential morphological operations from `scikit-image`: binary closing (`morphology.binary_closing, structure = disk(4)`), binary opening (`morphology.binary_opening, structure = disk(3)`), and binary dilation (`morphology.binary_dilation, structure = disk(3)`). Closing was applied first to fill small holes and protect tissue during opening. Opening removed small artefacts. Dilation added a conservative margin to reduce the risk that borderline tissue—often partially removed by the initial Otsu step—would be incorporated into the background class during background estimation. For very small tissue fragments at risk of being removed by opening, the opening structuring element was set to `disk(0)`.

Background mean intensity and standard deviation were estimated from pixels labelled as background. To address large slide-width artefacts, an artefact check was performed: if any elongated bright object spanned >90% of the slide width, the corresponding pixels were replaced by the estimated background mean, and the Stage 1 procedure (thresholding, morphology and background estimation) was repeated until no such artefact remained.

Because dilation purposedly overestimates the tissue region, a border refinement step was applied to remove misclassified background pixels at the tissue boundary: pixels within the tissue mask with intensity below (background mean + 3×SD) were reclassified as background. Using the refined tissue mask and the estimated background mean, a synthetic-background reconstructed grayscale image was generated by replacing background pixels with a uniform intensity equal to the background mean.

*Stage 2.* Triangle thresholding (`filters.threshold_triangle`) was applied to the synthetic-background reconstructed image. The resulting mask was then subjected to a light binary opening (`morphology.binary_opening, structure = disk(1)`) to remove residual small artefacts, followed by a final border refinement step using the same criterion (background mean + 3×SD) to sharpen tissue boundaries.

*Fatty tissues: combined hue–grayscale segmentation (Fig. M1b)*

For slides containing extensive fatty regions, grayscale intensity alone could not reliably separate tissue from background. Therefore, tissue detection primarily relied on the hue channel from low-resolution images converted to HSV space, while grayscale intensity was retained as a complementary cue because certain artefacts (and some tissue subregions) are more distinguishable in grayscale than in hue. In addition, fatty tissues frequently include detached small fragments and exhibit reduced contrast relative to background; consequently, these slides are more sensitive to background-based border refinement and to opening operations, and the pipeline prioritizes tissue preservation when required.

*Stage 1.* To preserve faint fatty tissue whose hue values were only marginally different from the background, local thresholding was used as the initial detection step: `filters.rank.otsu` on the hue channel with `footprint = disk(1)`. The resulting candidate tissue mask was refined by binary opening (`morphology.binary_opening`, `structure=disk(3)`) and binary dilation (`morphology.binary_dilation, structure=disk(3)`). This local-threshold-derived mask maximizes tissue retention but may also retain background artefacts and was therefore not used for background statistics estimation.

To ensure robust background estimation, a separate stringent background mask was generated from the hue channel using global Otsu thresholding (`filters.threshold_otsu`) followed by binary closing (`morphology.binary_closing`, `structure = disk(4)`). In rare cases where closing caused the entire slide to be labelled as foreground, closing was omitted, and global Otsu was applied alone. This stringent mask was used exclusively for estimating background statistics: background mean, and standard deviation were computed for both grayscale intensity and hue values.

Using these background estimates, synthetic-background reconstructed hue and grayscale images were generated (background replaced by the corresponding background mean), and slide-length artefact detection/correction was applied as described for non-fatty tissues. Border refinement was performed in two steps: first on hue using (background hue mean + 1.5×SD), then on grayscale using (background grayscale mean + 1.5×SD). The reduced multiplier (1.5 rather than 3) was used because fatty tissue often lies close to background values, and stricter thresholds would remove true tissue.

At this stage, the tissue candidate mask may still underestimate tissue extent. To add a conservative safety margin, we applied the following morphological sequence: binary closing (`disk(4)`), binary dilation (`disk(7)`), binary closing (`disk(5)`), and binary opening (`disk(3)`). A synthetic-background hue image was reconstructed using this expanded mask, and global Otsu thresholding on hue followed by binary closing (`disk(4)`) produced a tissue-plus-margin mask for subsequent refinement. Synthetic-background grayscale and hue images were regenerated on this mask.

*Stage 2.* To refine fatty tissue masks while limiting tissue loss, we combined grayscale and hue information in two steps: First, intersection refinement: the intersection of (i) Triangle thresholding (`filters.threshold_triangle`) on the synthetic-background grayscale image followed by binary closing (`morphology.binary_closing, disk(5)`) and (ii) local Otsu thresholding (`filters.rank.otsu, footprint=disk(1)`) on the synthetic-background hue image. Second, union refinement: synthetic-background grayscale and hue images were regenerated from the intersection mask, and the final mask was defined as the union of (i) Triangle thresholding on the reconstructed grayscale image and (ii) global Otsu thresholding (`filters.threshold_otsu`) applied to a hue image smoothed with a local mean filter (`filters.rank.mean, disk(1)`).

Finally, a light binary opening (morphology.binary_opening, structure = disk(2)) was applied to remove small residual artefacts introduced by the preceding operations.

### 2.3 WSI tiling

Segmented images were tiled into 224 × 224 pixel^2^ patches with `deepzoom.DeepZoomGenerator` from the `OpenSlide` library version 1.2.0. Tiles containing more than 50% background pixels were discarded. Background thresholds were determined for each WSI at the tissue segmentation step.

### 2.4 Tile embedding

Lunit-DINO^2^ was selected as our embedding model. This model, with a ViT-S backbone (22 million parameters), was pretrained on 19 million tiles from 15,672 WSI from TCGA and was evaluated on multiple pathology benchmarks (BACH, CRC, MHIST, PatchCamelyon, CoNSeP). A recent benchmark showed that its performances are not far off those of much larger models in a range of tiles- and WSI-level classification tasks despite its far superior computational efficiency, making it an appropriate choice for large-scale analyses^3^.

### 2.5 Morphemes identification

Morpheme identification proceeded independently for each organ. Embedding vectors were clustered into 64 morphemes using a Gaussian Mixture Model with `mixture.GaussianMixture`, from the `scikit-learn` library version 1.2.2, with parameters `n_components` = 64, `covariance_type` = `’diag’`, `n_init` = 10, and `random_state` = 0.

Morphemes were visually inspected, those containing tiles with blur, bubbles, fold and other technical effects were flagged as artefacts and removed from organ-specific morpheme count matrices (Table 1). Artefacts morphemes were excluded from all computations, but can be examined in the online atlas.

**Table 1.** Morphemes and slides discarded after atlas generation. Off target slides not reported by GTEx pathologists and artefacts-related morphemes were identified during our analyses and excluded from morpheme count preprocessing and subsequent analyses (Supp. Table S8). More slides have been discarded before atlas generation (Section 2.1).

| **Organ** | **Artifactual morphemes** | **Blacklisted slides** |
|---|---|---|
| AdiposeTissue | 2, 4, 56 | — |
| Aorta | 30 | — |
| AtrialAppendage | 28, 61 | — |
| Colon | 57 | GTEX-11WQK, GTEX-YB5K |
| EsophagusMucosa | 53 | — |
| EsophagusMuscularis | 49 | — |
| GastroesophagealJunction | 56 | GTEX-1HBPN |
| Heart | 56 | — |
| Ileum | 13, 55 | — |
| KidneyCortex | 26 | GTEX-139TT |
| Liver | 62 | — |
| Lung | 26, 31 | — |
| MammaryTissueBreast | 9, 57, 63, 50 | — |
| Omentum | 8, 12, 37, 56 | — |
| Pancreas | 52, 34 | GTEX-U3ZN, GTEX-U4B1, GTEX-U3ZM, GTEX-U3ZH, GTEX-1S3DN |
| SigmoidColon | 41 | GTEX-11WQK |
| SkeletalMuscle | 59, 16 | GTEX-O5YV |
| Skin | 43 | — |
| Stomach | 29, 33 | GTEX-05YV |
| SuprapubicSkin | 53 | — |
| ThyroidGland | 60 | — |
| TibialArtery | 45, 56 | GTEX-11ZTS, GTEX-11WQK |
| CoronaryArtery | 56, 21 | — |
| AdrenalGlands | 49 | — |
| Ovary | 48 | — |
| Uterus | 57 | — |
| Vagina | 26, 10, 30 | — |
| Spleen | 49, 54 | GTEX-PWOO |
| UrinaryBladder | 17, 47 | — |
| PituitaryGland | 36 | — |
| BrainCortex | 24, 26, 33 | GTEX-13OW8 |
| BrainCerebellum | 40 | — |
| Prostate | 56 | — |
| Testis | 22, 38, 40, 61 | GTEX-144FL |
| TibialNerve | 7, 24, 63 | GTEX-145MN, GTEX-1RB15, GTEX-11ZTS, GTEX-OHPJ, GTEX-1K2DU, GTEX-11WQK |
| MinorSalivaryGlands | 61 | — |

### 2.6 Morpheme quantification

Given an atlas of *K* morphemes, the raw expression of morpheme *j*, in a WSI, *i*, made of a set of tiles *T_i_* is defined as

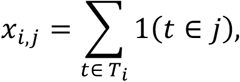

where 1(·) is the indicator function: 1(*x*) = 1 if *x* is true and 1(*x*) = 0 otherwise. The raw morphome of *i*, is defined as

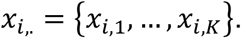

We set *K* to 64 (Section **Error! Reference source not found.**).

We encountered a few pairs of slides targeting the same organ in the same donor and summed their raw morphomes. We also noticed off target slides not labeled ‘Unacceptable’ during our biological investigations and removed them from the raw count matrices (Table 1).

Raw morphemes counts were next transformed to prevent the unwanted effects of massive zero-inflation and compositional structure, i.e. the fact that large expression of a morpheme implies lower expression of other morphemes. The per-organ, per-morpheme distribution of zeros are shown in Fig. M2a. The steps of the preprocessing and their effect on count distribution are illustrated Fig. M2b. We first normalized morpheme counts as to set their slide-wise sum to 1. To prevent computational issues related to the zero-inflation of morphemes counts, we applied multiplicative replacement, a standard method in compositional data analysis^4^. The zero-valued morphemes values, whose indices were *Z_i_* = {*j*: *x_i_*_j_ = 0}, were replaced with:

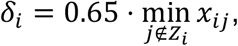

while non-zero values are proportionally adjusted by a factor,

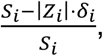

to preserve the sum of morpheme counts equal to 1. We used the default replacement factor, 0.65, from ref. ^4^.

Because of the compositional nature of the data, morphemes lie on a *K* − 1 simplex, which induces spurious negative correlations between morphemes. To address this, we applied the centered log-ratio transformation^5^,

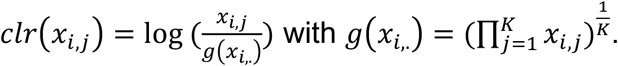

Finally, to fulfill constant variance of the morphemes for all downstream association tasks (those including confounders-adjustments), we employed inverse normal transformation:

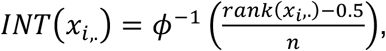

where *φ*^−1^ denotes the quantile function of the standard normal distribution and *rank*(*x_i_*) is the average rank of observation *x_i_* normalized in [0,1] among all non-discarded samples. Ties were handled by assigning average ranks. This rank-based transformation maps the data to a standard normal distribution, ensuring homoscedasticity and robustness to outliers while preserving the relative order of observations. The cross-organ, cross-morphemes densities at the final step are shown in Fig. M2c.

**Fig. M2.**
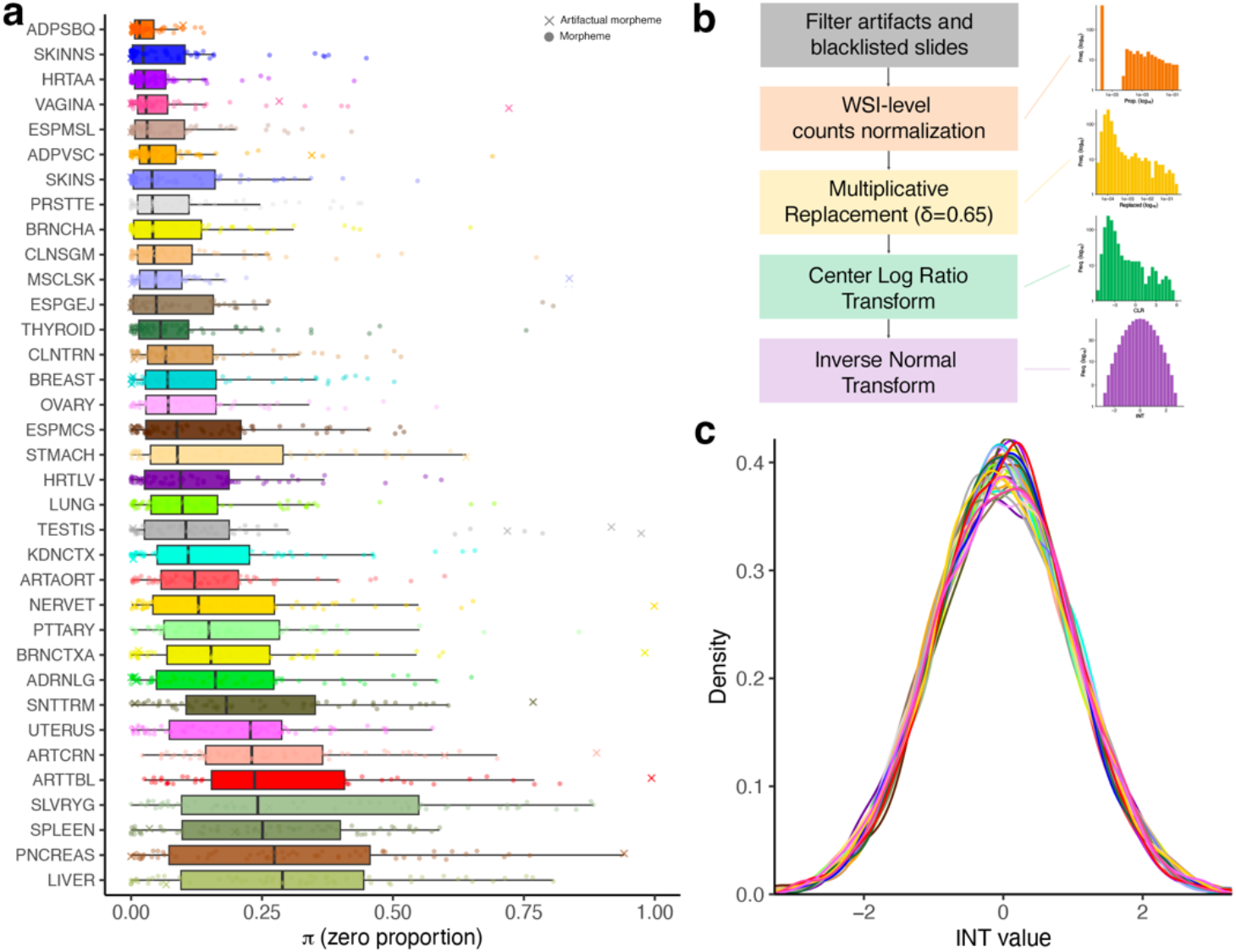
Preprocessing of morpheme raw counts. a, Per-organ distributions of zeros. Each dot represents a morpheme; crosses denote morphemes that were identified as artefactual post-hoc. **b**, The 5 steps of morphemes pre-processing and their effect on morphemes’ distributions are illustrated: first, artefact morphemes and blacklisted slides were filtered out; second, for a given organ, the morphome of each WSI was normalized by the total number of tiles it contained. Third, to attenuate zero-inflation, multiplicative replacement was applied to each WSI. Fourth, a centered log-ratio transformation was applied to handle data compositionality. Fifth, expression values were inverse normal-transformed to enforce normality. **c**, The final distributions were normally distributed (each line represents a morpheme within an organ). This pre-processing pipeline is key to ensure that statistical associations are not biased by skews and zero-inflation.

### 2.7 Correlation analyses

Molecular data were preprocessed according to modality-specific requirements:

- Inverse-normal transformed telomere length (a.k.a. Telomere Quantity Index, `TQImean`) and protein abundance data were downloaded from the GTEx portal. No further processing was applied.
- Small RNA counts, restricted to miRNA and TPM-normalized were downloaded from the GTEx portal (filename miRNA_TPM_matrix_PORTAL_2025_03_17.txt.gz). For differential expression, TPMs were (log(.)+1)-transformed.
- For methylation data, the 20,000 most variable CpG probes were selected, β-values were converted to M-values using the logit transformation 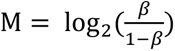
- We downloaded from the GTEx portal pathology categories derived from free text pathology notes by the GTEx consortium. Categories were considered for correlation analyses and classification tasks if at least 15 cases and 30 cases were available, respectively.

For long and small RNA transcriptomes, differential expression analysis was performed using `edgeR`^6^ version 3.36.0 with quasi-likelihood generalized linear models and TMM normalization. Genes were filtered using the function `filterByExpr` with default parameters (`min.count` = 10, `min.total.count` = 15, `min.prop` = 0.7).

For all other molecular modalities (proteins, telomeres, small RNA, and methylation), `limma`^7^ version 3.50.0 linear models were used to test associations between molecular features and morpheme expression. All models were adjusted for age, sex, ischemic time, and cohort, with sex excluded from models for sex-specific organs. We applied the Benjamini-Hochberg method to control the false discovery rate.

### 2.8 Transcriptional signatures enrichment, including cell type signatures

To interpret the biological significance of morpheme expression, we performed gene set enrichment analysis using cell type-specific markers from the *PanglaoDB*^8^ database. We first filtered markers to retain only those labeled in *PanglaoDB* as ‘canonical’ from the RNA-seq transcriptomes of the organs of interest. Next, genes were ranked by signed significance statistic from the differential expression analysis performed with `edgeR` as described above, i.e. genes were ranked according to *rank*[− log_10_(*FDR*) ⋅ *sign*(*FC*)]). Finally, enrichments were calculated using the R package `fgsea`^9^ version 1.20.0. This approach enabled quantification of whether specific morphological clusters were associated with enrichment of cell types characteristic of the organ of interest. Organ-specific cell types were combined with cell types from ubiquitous tissues such as vasculature, blood, connective tissue, epithelium, immune system and muscle. Results are reported as normalized enrichment scores (NES) and associated FDR.

In the analysis of main text Fig. 5f, we used HALLMARK_G2M_CHECKPOINT signature from MSigDB^10^ and the positive markers of papillary thyroid cancer listed in ref.^11^.

### 2.9 Adipocyte segmentation and association with molecular adipocyte signature

Tile-level adipocyte content was quantified using `scikit-image`^12^ version 0.19.3 by applying grayscale thresholding to identify adipocyte regions (intensity > 225), followed by morphological cleaning (hole removal, binary closing with disk radius 3, and binary opening with disk radius 2) to remove small artefacts. Connected components smaller than 1,000 pixels were excluded, and the proportion of adipocyte area per tile was calculated from the resulting binary mask.

To compute the association between difference between M**11** and M**27** (Fig. 4f in main text), canonical adipocyte marker genes were obtained from PanglaoDB^8^, and GSEA with the package `fgsea`^9^ was performed on genes ranked by log fold-change from differential expression analysis (adjusted for age, cohort, ischemic time, and BMI) to identify leading edge genes. The adipocyte score was computed as the sum of CPM-normalized expression with these leading edge genes.

### 2.10 Predictions from morphemes

Classification and regression tasks were conducted using the `scikit-learn`^12^ version 0.24.2. Pathology keywords were considered for classification task if a given pathology had more than 30 cases. Classification was performed using `LogisticRegression` with L2 penalty and `StratifiedKFold` cross-validation to maintain class proportions. Regression was performed using `RidgeCV` with standard `KFold` splitting. For both task types, we implemented nested cross-validation—an inner loop for tuning regularization strength and an outer `KFold`=5 loops for unbiased evaluation. To avoid data leakage, transformed morpheme expression was residualized against confounders using parameters derived exclusively from training data. Results were reported as `KFold`-averaged AUCs for classification tasks and `KFold`-averaged *R*^2^ for regression.

### 2.11 Predictions from whole blood transcriptomes

Whole blood long RNA raw counts were processed with `edgeR`^6^: We selected genes with a minimum count of 5 per sample, a minimum total count of 10 across samples with `filterByExpr`. Counts were then normalized to counts per million (CPM) without TMM normalization to preserve interpretable per-sample scaling appropriate for downstream machine learning applications.

Whole blood small RNA raw counts were filtered as to retain only annotated miRNAs by mapping gene identifiers to miRbase^13^ names using curated annotations. When multiple identifiers mapped to the same miRNA, expression values were averaged, and the resulting counts were normalized to CPM.

For morpheme prediction tasks, continuous morpheme values were binarized at a threshold treated as a hyperparameter optimized through nested cross-validation. Classification was performed using `LogisticRegression` with Elastic Net regularization and balanced class weights. Nested cross-validation included an inner 3-fold loop tuning the regularization strength (`C`), the L1/L2 mixing ratio (`l1_ratio`), and the morpheme percentile-based binarization threshold. An outer 5-fold cross-validation loop provided unbiased performance estimates. To prevent data leakage, gene selection (top 2,500 most variable genes by coefficient of variation), threshold computation, residualization of gene expression and morpheme values against confounders (age, sex, cohort and ischemic time), and feature scaling were all performed independently within each outer fold using only training data. Feature importances were aggregated across folds rather than derived from a single model fitted on all data.

### 2.12 Penalty setting in ML models

Elastic Net regularization was used when predicting from transcriptomes, which contain thousands of features, whereas Ridge regression was used when predicting from morphomes, containing up to 64 morphemes. Elastic Net combines L1 and L2 penalties, promoting sparsity to identify relevant features while maintaining stability when predictors are correlated, which is often the case with transcriptomic data.

### 2.13 Cross-organ morpheme expression predictions

To quantify shared histological signals across tissues, we assessed how well morpheme expression in each organ predicted morpheme expression in every other organ. Morpheme values were first residualized against ischemic time, cohort, sex, and age. For each target morpheme, we fit a linear model using all morphemes from a predictor tissue as features and computed adjusted R². Results were summarized by computing the median adjusted R² across all morphemes for each tissue pair, yielding a tissue × tissue predictability matrix. Hierarchical clustering used correlation distance and Ward linkage.

### 2.14 Genome-wide association studies

We followed the trans eQTL discovery procedure of the GTEx consortium^14^, with the following variations. First, we started from morpheme counts preprocessed as described Section 2.6. Second, we upgraded genomes vcf files from GTEx v8 to v10 following the GTEx consortium procedures^14^. Third, we computed associations with `tensorqtl`^15^, a GPU-accelerated implementation of the `fastqtl` algorithm used by the consortium. Fourth, we included the following covariates in the model: age, sex, cohort (postmortem/organ donor+surgical), genotyping platform, PCR protocol, ischemic time, and the first 5 genotype principal components available in the GTEx portal. Variants with minor allele frequency (MAF) <0.05 were excluded (∼6.5 million variants remained, Suppl. Fig. S1b).

Genome-wide significance threshold proposed as guide in the atlas are genome-wide significance, p<5×10^-8^; organ-wise significance, p<7.81×10^-10^ (p<5×10^-8^/64) and atlas-wise significance at p<1.95×10^-11^ (p<5×10^-8^/(64×40)).

### 2.15 GWAS colocalization analyses

Colocalizations between hQTL signals, GTEx v10 RNA expression QTLs and GWAS signals from the GWAS catalog were performed using the Bayesian approximate Bayes factor method implemented in the `coloc.abf` function of the R package `coloc`^16^, which assumes a single causal variant per locus for each trait. For each hQTL lead variant. Variants within a ±500 kb window were considered. GTEx eQTL full summary statistics were pre-filtered by molecular phenotype. We use the harmonized versions of studies from the GWAS Catalog^17^, full statistics were used. Default prior probabilities were applied.

### 2.16 Thyroid epithelial morphemes and junction protein score

To assess epithelial integrity, we defined a tight junction protein signature comprising 12 genes encoding claudins (CLDN1, CLDN3, CLDN4, CLDN5), occludin (OCLN), zonula occludens proteins (TJP1, TJP2), junctional adhesion molecule (F11R), and adherens junction components (CDH1, CTNNB1, CTNNA1, CTNND1); based on the inverse-normal transformed mean expression. We then compute the difference between thyroid M**63** and M**47** (Suppl. Fig. 6), and a linear regression is fitted without additional covariates:

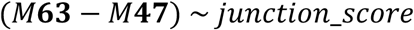

### 2.17 Consensus minimum spanning tree construction

To characterize morpheme relationships in the morphological space, we constructed a bootstrap consensus minimum spanning tree (MST). For each of *n*=100 bootstrap iterations, we subsampled up to 2,000 tiles per morphemes and computed their centroids as their mean embedding vector. Pairwise Euclidean distances between centroids were computed using `scipy.spatial.distance.cdist`, and an MST was extracted via `networkx.minimum_spanning_tree` from the `networkx` library^18^ version 2.6.3. The consensus MST retained edges appearing in ≥40% of bootstrap iterations, with edge frequency quantifying topological stability. Representative tile paths between adjacent morphemes were identified by applying `scipy.sparse.csgraph.shortest_path` from `scipy`^19^ version 1.6.0 on a *k*-nearest-neighbor graph (*k* =50) constructed using `cuml.neighbors.NearestNeighbors` from `cuml`^20^ version 22.12.0.

### 2.18 Validation of kidney cortex morphemes blood transcriptome signatures in external uremia dataset

To validate kidney cortex morpheme-derived gene signatures, we used the GSE37171 dataset from GEO comprising peripheral blood transcriptomes from 75 patients with end-stage renal disease and 40 healthy controls, profiled on Affymetrix Human Genome U133 Plus 2.0 arrays^21^. For morphemes M**35** and M**39**, genes selected across all five cross-validation folds of each morpheme were retained. We focused on *TPSAB1* and *TPSB2*, which encode tryptase subunits characteristic of mast cells, as a tryptase signature. For each sample in GSE37171, individual gene expression values were z-score-normalized across samples, and the signature score was calculated as the mean of the normalized expression values for *TPSAB1* and *TPSB2*, followed by inverse-normal transformation and residualization for the sex, age, and ‘race’ covariates reported in GSE37171. Discriminative performance for the ‘healthy’ versus ‘uremia’ comparison was assessed using area under the ROC curve (AUC) via `roc` from `pROC`^22^ package version 1.18.0. Confidence intervals were estimated by stratified bootstrap resampling (n = 1,000). To control for the biological specificity of the validation AUC obtained for *TPSAB1* and *TPSB2,* we ran the entire validation protocol on 1,000 pairs of genes selected randomly in the human genome—a control inspired from ref. ^23^. We computed specificity as the fraction of random pairs giving lower AUC than the pair *TPSAB1*:*TPSB2*.

